# Kap-β2/Transportin mediates β-catenin nuclear transport in Wnt signaling

**DOI:** 10.1101/2021.05.22.445277

**Authors:** Woong Y. Hwang, Valentyna Kostiuk, Delfina P. González, C. Patrick Lusk, Mustafa K. Khokha

**Author notes:** correspondence to: C. Patrick Lusk, Mustafa K. Khokha.

## Abstract

Wnt signaling is essential for many aspects of embryonic development including the formation of the primary embryonic axis. In addition, excessive Wnt signaling drives multiple diseases including cancer highlighting its importance for disease pathogenesis. β-catenin is a key effector in this pathway that translocates into the nucleus and activates Wnt responsive genes. However, due to our lack of understanding of β-catenin nuclear transport, therapeutic modulation of Wnt signaling has been challenging. Here, we took an unconventional approach to address this long-standing question by exploiting a heterologous model system, the budding yeast *Saccharomyces cerevisiae*, which contains a conserved nuclear transport machinery. In contrast to prior work, we demonstrate that β-catenin accumulates in the nucleus in a Ran dependent manner, suggesting the use of a nuclear transport receptor (NTR). Indeed, a systematic and conditional inhibition of NTRs revealed that only Kap104, the orthologue of Kap-β2/Transportin-1 (TNPO1), was required for β-catenin nuclear import. We further demonstrate direct binding between TNPO1 and β-catenin that is mediated by a conserved amino acid sequence that resembles a PY NLS. Finally, using *Xenopus* secondary axis and TCF/LEF reporter assays, we demonstrate that our results in yeast can be directly translated to vertebrates. By elucidating the NLS in β-catenin and its cognate NTR, our study provides new therapeutic targets for a host of human diseases caused by excessive Wnt signaling. Indeed, we demonstrate that a small chimeric peptide designed to target TNPO1 can reduce Wnt signaling as a first step towards therapeutics.

## INTRODUCTION

Wnt signaling plays multiple roles in embryonic development. For example, Wnt signaling is critical for establishing the dorsal embryonic axis; increased Wnt signaling can lead to a secondary axis (twinning of the embryo) while depletion of a key effector of the Wnt signaling pathway, β-catenin, can lead to a radially ventralized embryo (Heasman et al., 1994; Heasman, Kofron, & Wylie, 2000; Khokha, Yeh, Grammer, & Harland, 2005; McMahon & Moon, 1989; Moon, Brown, & Torres, 1997; Nishisho et al., 1991; Smith & Harland, 1991; Sokol, Christian, Moon, & Melton, 1991). In addition, Wnt signaling has been implicated in a variety of human diseases especially cancer (Clevers & Nusse, 2012; MacDonald, Tamai, & He, 2009; Moon, Kohn, De Ferrari, & Kaykas, 2004; Morin et al., 1997; Nusse & Varmus, 1982; Polakis, 2012; Wood et al., 2007). In fact, 90% of colorectal cancers are caused by genetic alterations in Wnt pathway factors (Cancer Genome Atlas, 2012). Therefore, chemical inhibitors of Wnt signaling have tremendous therapeutic potential.

In Wnt signaling, β-catenin (CTNNB1) relays the message from a Wnt ligand at the plasma membrane to transcription factors in the nucleus (MacDonald et al., 2009; Niehrs, 2012). As such, its levels are kept under control through a constitutively active degradation pathway. In the presence of Wnt ligand, the degradation machinery is sequestered, and the resulting stabilization of β-catenin allows it to enter the nucleus where it drives the transcription of Wnt responsive genes (MacDonald et al., 2009; Niehrs, 2012). Despite intensive study, the mechanism of β-catenin translocation from the cytosol to the nucleus remains obscure. First, while early studies suggested that β-catenin nuclear import was energy dependent, they excluded a role for the Ran GTPase (Fagotto, Gluck, & Gumbiner, 1998; Yokoya, Imamoto, Tachibana, & Yoneda, 1999), the master regulator of the nuclear transport of most macromolecules bearing either nuclear localization signals (NLS) and/or nuclear export signals (NES) (Cavazza & Vernos, 2015). Second, prior studies demonstrated that β-catenin nuclear import did not require the Kap-α/Kap-β1 (Importin α/β1) nuclear transport receptor (NTR) complex (Fagotto et al., 1998; Yokoya et al., 1999), a notion consistent with Ran independence. Finally, the structural similarity between β-catenin and Kap-α, both made up of armadillo-repeats, suggested that β-catenin might itself act as an NTR by directly interacting with the Phe-Gly (FG) nups responsible for selective passage across the nuclear pore complex (NPC) (Andrade, Petosa, O’Donoghue, Muller, & Bork, 2001; Conti & Kuriyan, 2000; Fagotto et al., 1998; A. H. Huber, Nelson, & Weis, 1997; Sharma, Johnson, Brocardo, Jamieson, & Henderson, 2014; Xu & Massague, 2004; Yano, Oakes, Tabb, & Nomura, 1994). However, Kap-α does not directly bind to FG-nups and the evidence that β-catenin does so is controversial (Sharma, Jamieson, Lui, & Henderson, 2014; Suh & Gumbiner, 2003). Using a CRISPR based screening platform, recent work provides evidence that β-catenin may be imported by the NTR, Imp11; however, this mechanism appears curiously relevant only for a subset of colon cancer cells (Mis et al., 2020). Alternative models for β-catenin nuclear transport have been proposed; however, molecules that directly bind β-catenin to modulate nuclear transport remain undefined (Goto et al., 2013; Griffin et al., 2018; Komiya, Mandrekar, Sato, Dawid, & Habas, 2014).Thus, a complete understanding of β-catenin nuclear transport remains outstanding, leaving open a key gap in our knowledge of Wnt signaling that could otherwise be targeted for therapeutic intervention.

A major challenge with understanding the β-catenin nuclear import mechanism is the myriad of binding partners that modulate its steady-state distribution and, hence, complicate the direct interrogation of the nuclear transport step (Fagotto, 2013; MacDonald et al., 2009). Here, we take an unconventional approach and investigate β-catenin nuclear transport in a heterologous system, the budding yeast *S. cerevisiae*. Yeast do not have a Wnt pathway nor a β-catenin ortholog, which presumably emerged in metazoans with the onset of multicellularity and cell fate specialization (Holstein, 2012). Yeast also likely lack β-catenin binding partners and its degradation machinery and thus provide a simplified system to specifically evaluate nuclear import. Most critically, the nuclear transport system including NTRs, NPCs and Ran are well conserved from yeast to human (Malik, Eickbush, & Goldfarb, 1997; Wente & Rout, 2010; Wozniak, Rout, & Aitchison, 1998). Indeed, even NLS and NES sequences are recognized by orthologous NTRs across millions of years of evolution (Conti & Kuriyan, 2000; Fontes, Teh, & Kobe, 2000; Kosugi, Hasebe, Tomita, & Yanagawa, 2008; Lange, Mills, Devine, & Corbett, 2008; Soniat et al., 2013). Using the yeast system as a discovery platform, we uncover a PY-like NLS in B-catenin that can be directly recognized and imported into the nucleus of yeast, *Xenopus* and human cells by Kap-β2/Transportin-1 (TNPO1). We further demonstrate that a small peptide based on a chimeric PY-NLS sequence can prevent Wnt signaling, opening the door for therapeutics.

## RESULTS

### β-catenin requires a functional Ran GTPase to localize in the nucleus of *S. cerevisiae*

Investigating the β-catenin nuclear import mechanism in yeast relies on the premise that a minimal, conserved β-catenin transport machinery exists in this organism. Therefore, we first tested whether β-catenin accumulates in the yeast nucleus. Specifically, we assessed the localization of a *Xenopus* β-catenin-GFP (xβ-catenin-GFP) in a wildtype yeast strain expressing an endogenously tagged nuclear envelope membrane protein, Heh2-mCherry, to help visualize the nuclear boundary. Indeed, xβ-catenin-GFP was enriched in the nucleus compared to GFP alone (Figure 1A). To quantify this steady state distribution, we measured the nuclear enrichment of xβ-catenin-GFP by relating the mean GFP fluorescence in the nucleus (N) and cytoplasm (C) (Figure 1A, plot at right). The xβ-catenin-GFP had a mean N:C ratio of ∼1.6, which was significantly higher than GFP alone (1.1). As xβ-catenin-GFP is 119 kD, it would be unable to easily pass through the NPC diffusion barrier suggesting that xβ-catenin-GFP can access a facilitated nuclear transport mechanism through the NPC (Popken, Ghavami, Onck, Poolman, & Veenhoff, 2015; Timney et al., 2016; Weis, 2003).

**Figure 1.**
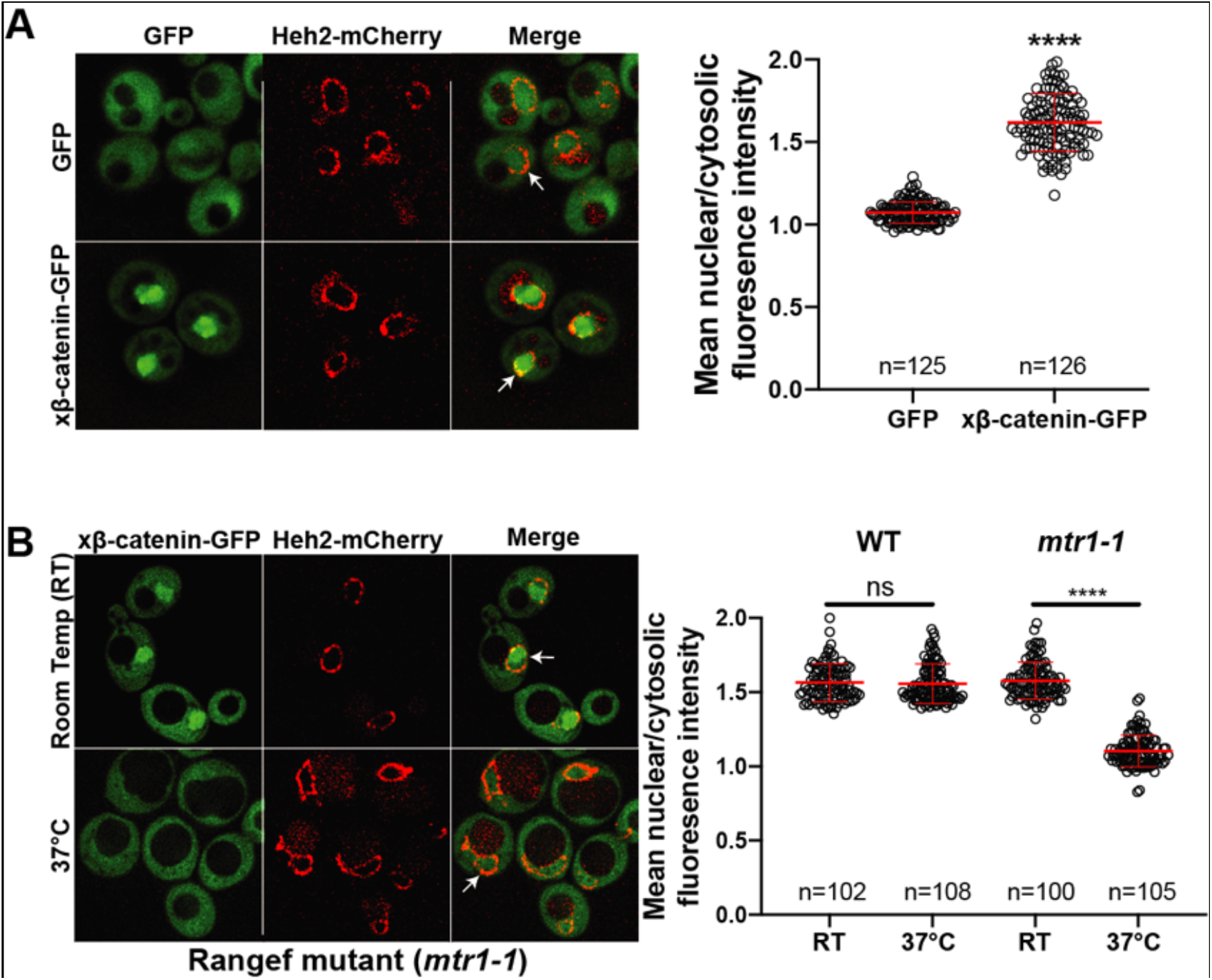
β-catenin requires a functional Ran GTPase to localize in the nucleus of *S. cerevisiae*. **(A)** Representative deconvolved fluorescence image of xβ-catenin-GFP in a wild-type yeast strain that expresses Heh2-mCherry to label the nucleus (left). White arrows indicate the nuclear compartment. Plot showing the quantification of mean nuclear to cytosolic fluorescence intensity from three independent replicates (right). **(B)** Representative deconvolved fluorescence image of xβ-catenin-GFP in the RanGEF mutant (*mtr1-1*) strain at room temperature or 37°C that co-expresses Heh2-mCherry as a nuclear envelope marker (left). The ratio of mean nuclear to cytosolic fluorescence intensity was measured in the wild-type or *mtr1-1* strain from three independent replicates (right). Red bar indicates the mean value with the standard deviation (SD). *p*-values are from unpaired two-tailed t-test where ns is p>0.05, and ****p<0.0001 for both **(A)** and **(B)**.

In principle, three mechanisms of xβ-catenin nuclear import are possible: 1) xβ-catenin-GFP is imported by an NTR, 2) xβ-catenin-GFP piggybacks on an unknown binding partner that is itself imported by an NTR or 3) xβ-catenin-GFP has an intrinsic ability to cross the NPC free of NTRs. To rule out the latter possibility, we tested whether xβ-catenin-GFP nuclear accumulation was dependent on a functional Ran gradient, which would specifically impact NTR-mediated transport (Schmidt & Gorlich, 2016; Weis, 2003; Wente & Rout, 2010). We therefore assessed β-catenin-GFP localization in the *mtr1-1* mutant strain, which is a temperature sensitive, loss of function allele in the gene encoding the yeast Ran-GEF (*SRM1*/*MTR1/PRP20*) (Kadowaki, Zhao, & Tartakoff, 1992). As Ran-GEF exchanges GDP for GTP on Ran in the nucleus, it is essential for the functioning of the nuclear transport system (Weis, 2003). At room temperature, xβ-catenin-GFP is localized in the nucleus with a N:C ratio similar to the wild type strain (∼1.6) (Figure 1B). In striking contrast, growth at 37°C, which is non-permissive for Mtr1p function, results in the re-distribution of xβ-catenin-GFP such that it is evenly distributed between the nucleus and cytoplasm with N:C ratios identical to GFP alone (1.1) (Figure 1B). Importantly, xβ-catenin-GFP localization was not affected by the elevated temperature as wild-type cells showed N:C ratios of ∼1.6 even at 37°C. Thus, nuclear accumulation of xβ-catenin-GFP is dependent on a functioning Ran GTPase system raising the possibility that it requires a NTR-mediated pathway to accumulate in the nucleus.

### The C-terminus of β-catenin contains a NLS

Having established that β-catenin import requires a functional Ran pathway, we next sought to map the sequence elements of β-catenin that confer nuclear localization. β-catenin can be divided into three domains: a central region rich in armadillo (ARM) repeats sandwiched between two unstructured domains (Figure 2A). We generated constructs where each of these domains was individually deleted and examined their localization in yeast. Removal of the C-terminus significantly reduced nuclear enrichment of xβ-catenin-(1-664)-GFP (mean N:C values of 1.4) compared to constructs lacking either the N [xβ-catenin-(141-782)-GFP] or ARM (Δ141-664) domains, which accumulated in the nucleus at levels similar to the full length protein (mean N:C of 1.6) (Figures 2A-C). These data suggested that the C-terminus contains sequence elements required for nuclear accumulation. Consistent with this idea, the C-terminus of β-catenin [xβ-catenin-(665-782)-GFP] was sufficient to confer nuclear accumulation of GFP to levels comparable to the full length protein (mean N:C 1.6) (Figures 2A, 2B, and 2D). Of note, both the ARM repeats [xβ-catenin-(141-664)-GFP]) and the N-terminus of β-catenin [xβ-catenin-(1-141)-GFP] could confer some nuclear localization of GFP but to a considerably lesser extent than the C-terminus (mean N:C of ∼1.3) (Figures 2A, 2B, and 2D). Additionally, the ARM repeats had some affinity for the nuclear periphery (Figures 2D, xβ-catenin-(141-664)-GFP, white arrows). Overall, while there may be several elements of xβ-catenin that, in isolation, can target to the nucleus, the C-terminus contains a sequence that was both necessary and sufficient for nuclear accumulation at levels comparable to the full length protein. Consistent with previous work (Koike et al., 2004; Mis et al., 2020), these data suggested that the C-terminus of xβ-catenin contained an NLS, which we further mapped to amino acids 665-745 (Figures 2A, 2B, and 2E).

**Figure 2.**
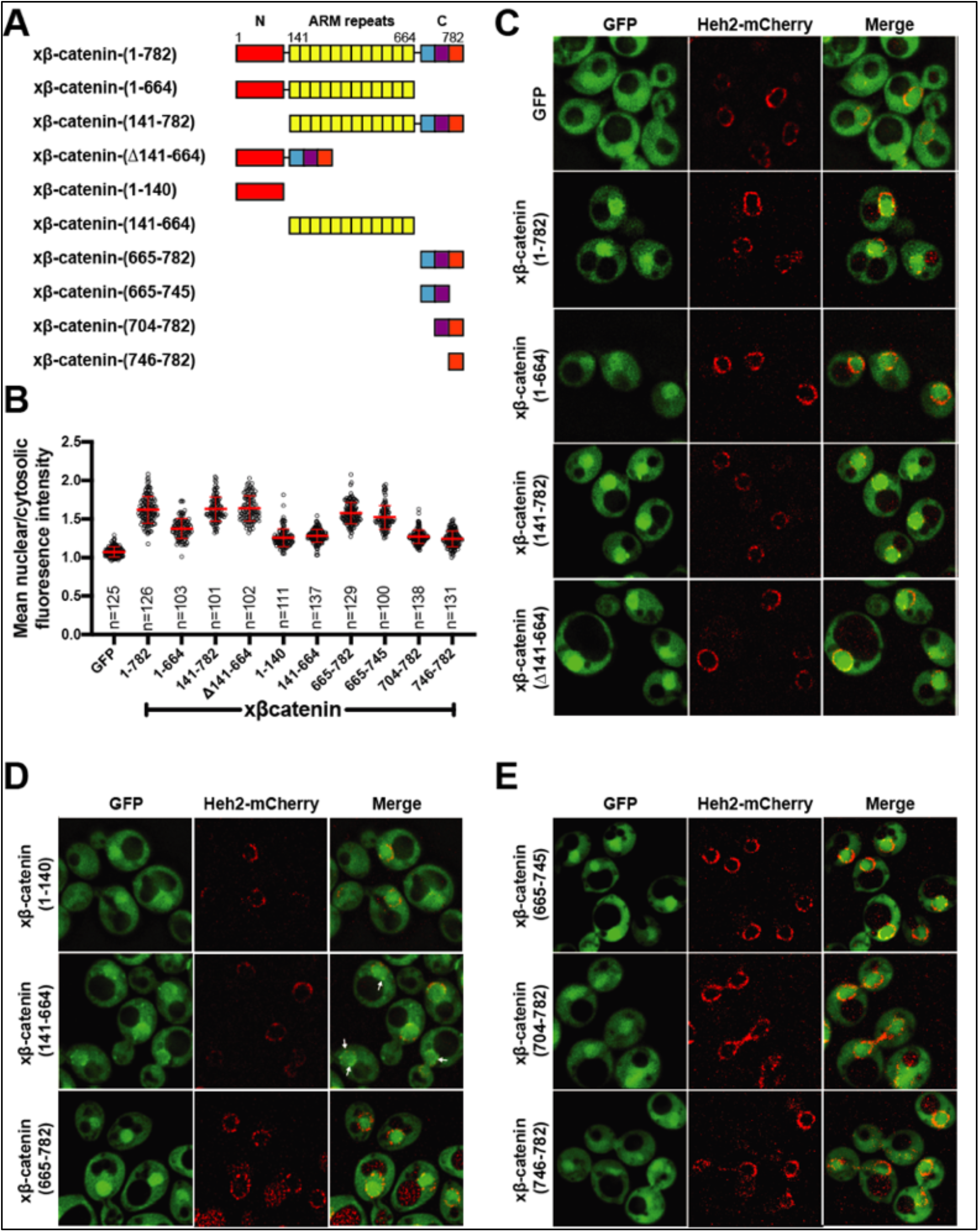
The C-terminus of β-catenin contains a NLS. **(A)** Schematic diagram of *Xenopus* β-catenin truncation constructs tested in this study. **(B)** Plot of the ratio of mean nuclear to cytosolic fluorescence intensity of *Xenopus* β-catenin GFP truncation constructs tested in a wild-type yeast strain from three independent replicates. Red bar indicates the mean value with the SD. **(C)** Deconvolved fluorescence image of the N-terminal deletion (141-782), ARM-repeats deletion (Δ141-664) or C-terminal deletion (1-664) of *Xenopus* β-catenin GFP in the wild-type strain. GFP and full-length *Xenopus* β-catenin-GFP were used as controls. **(D)** Deconvolved fluorescence images of the indicated fragments of *Xenopus* β-catenin GFP in the wild-type strain. White arrows indicate nuclear rim localization. **(E)** Deconvolved fluorescence images of indicated C-terminus fragments of *Xenopus* β-catenin GFP in the wild-type strain. Heh2-mCherry was co-expressed to label the nuclear membrane in **(C), (D)**, and **(E)**.

We confirmed that nuclear accumulation of xβ-catenin-(665-745)-GFP was dependent on the Ran pathway using the *mtr1-1* strain (Figure 2-figure supplement 1A). To ensure that this sequence did not confer binding to a yeast-specific factor, we also tested localization of xβ-catenin-(665-745)-GFP in HEK293T cells, a human embryonic kidney cell line. In line with the yeast results, xβ-catenin-(665-745)-GFP showed higher levels of nuclear accumulation compared to GFP alone (Figure 2-figure supplement 1B). Next, we investigated the function of the β-catenin NLS in the context of Wnt signaling using the secondary axis assay in *Xenopus*(McMahon & Moon, 1989; Smith & Harland, 1991; Sokol et al., 1991). Overexpression of Wnt effectors including β-catenin induces a secondary axis in *Xenopus* embryos. By injecting a moderate dose (200 pg) of xβ-catenin mRNA, secondary axes develop in roughly half of the embryos (Figure 2-figure supplement 2A-B) compared to none in the uninjected controls (UIC). If we delete the coding sequence for the NLS [xβ-catenin-(Δ665-745)-GFP], then the number of embryos with secondary axes is significantly reduced (Figure 2-figure supplement 2A-B). To ensure that this loss of function is due to the inhibition of β-catenin nuclear import, we added the classical NLS of the SV40 Large T-antigen (cNLS), which is imported by Kap-α/β1, to the N-terminus of β-catenin [cNLS-xβ-catenin-(Δ665-745)-GFP]. cNLS-xβ-catenin-(Δ665-745)-GFP could induce secondary axes similarly to the full length β-catenin (Figure 2-figure supplement 2B), consistent with the conclusion that xβ-catenin-(Δ665-745)-GFP was still functional for Wnt signaling but lacked the nuclear localization element. Supporting this supposition, we imaged xβ-catenin-GFP in these *Xenopus* embryos and found that while xβ-catenin-GFP localizes to the cell membrane and nucleus, xβ-catenin-(Δ665-745)-GFP shows a reduction in nuclear accumulation, which is rescued by the addition of the cNLS (Figure 2-figure supplement 2C).

### Kap104 is specifically required for β-catenin nuclear localization in *S. cerevisiae*

Next, to define the NTR responsible for xβ-catenin-GFP nuclear import, we used the Anchor-Away approach (Haruki, Nishikawa, & Laemmli, 2008) to systematically inhibit 10 budding yeast NTRs, all of which have orthologues in human cells (Table 1). This strategy takes advantage of the rapamycin-induced dimerization of a FK506 binding protein (FKBP12) with the FKBP-rapamycin binding (FRB) domain (Figure 3A). In this system, NTR-FRB fusions are expressed in a strain harboring FKBP12 fused to a highly abundant plasma membrane protein (Pma1) (Figure 3A). The addition of rapamycin leads to the rapid (∼15 min) trapping of the NTRs at the plasma membrane (Haruki et al., 2008). We systematically tested whether the addition of rapamycin (or the DMSO carrier alone) impacted the nuclear accumulation of xβ-catenin-(665-782)-GFP in each of the 10 NTR-FRB strains. Consistent with prior data (Fagotto et al., 1998), plasma membrane trapping of the Kapβ1 orthologue, Kap95-FRB, did not impact nuclear localization of xβ-catenin-(665-782)-GFP (Figure 3B). Indeed, trapping 9 of the 10 NTRs, including the Imp-11 orthologue, Kap120, had no overt influence on xβ-catenin-(665-782)-GFP nuclear localization (Figure 3B, 3C, and Figure 3– figure supplement 1). In contrast, we observed a remarkable inhibition of nuclear accumulation specifically when Kap104-FRB [orthologue of Kapβ2/Transportin-1(TNPO1)] was anchored away (Figure 3B and 3C). These data support a model in which Kap104 specifically mediates the nuclear import of xβ-catenin-(665-782)-GFP in the yeast system.

**Table 1.** List of human NTR genes and Yeast orthologs.

| Human NTR | Yeast Orthologs |
| --- | --- |
| Kap β1 | Kap95 |
| Kap $\alpha$ | Kap60/Srp1 |
| Transportin 1 | Kap104 |
| Importin-5/Kap β3 | Kap121/Pse1 |
| Importin-4/RanBP5 | Kap123 |
| Importin-7/RanBP7 | Kap119/Nmd5 |
| Importin-8/RanBP8 | Kap108/Sxm1 |
| Importin-9 | Kap114 |
| Importin-11 | Kap120 |
| Transportin-SR/TNPO3 | Kap111/Mtr10 |
| Importin-13 | Kap122/Pdr6 |
| CRM1/Exportin-1 | Xpo1 |
| Exportin-t | Los1 |
| Exportin-5 | Kap142/Msn5 |
| CAS | Cse1 |

**Figure 3.**
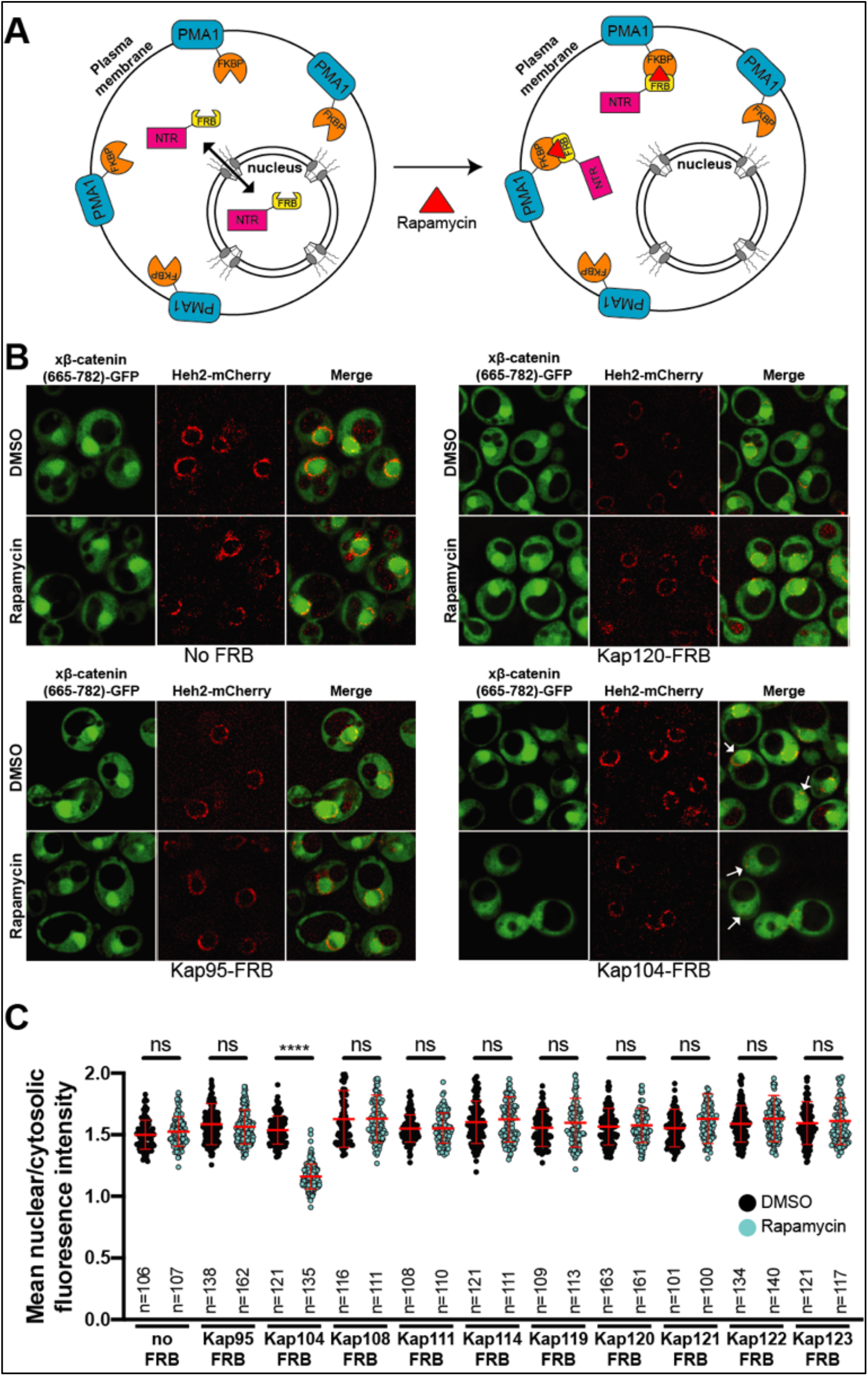
Kap104 is specifically required for β-catenin nuclear localization in *S. cerevisiae*. **(A)** Schematic of the Anchor Away assay mediated by the rapamycin induced dimerization of NTR-FRB and Pma1-FKBP12. Pma1 is a plasma membrane ATPase. **(B)** Deconvolved fluorescence image of Xenopus β-catenin (665-782)-GFP treated with DMSO (vehicle) or rapamycin for 15 min in the no FRB, Kap95 (Karyopherin β1 in human)-FRB, Kap120 (importin 11 in human) and Kap104 (Kapβ2/Transportin 1 in human)-FRB. Heh2-mCherry was used as a nuclear envelope marker. White arrows indicate the nuclear compartment **(C)** Plot showing the ratio of mean nuclear to cytosolic fluorescence intensity of Xenopus β-catenin (665-782)-GFP in the 10 NTR-FRB strains treated with DMSO or rapamycin. Red bar indicates the mean value with the SD. Experiments were performed three times. p-values are from unpaired two-tailed t-test where ns is p>0.05, and ****p<0.0001.

### β-catenin contains a PY-like NLS that is required for direct binding to TNPO1

Having established that Kap104 mediates β-catenin nuclear transport in yeast, we compared the xβ-catenin-(665-782) protein sequence to established TNPO1 NLS sequences (Lee et al., 2006; Soniat & Chook, 2015; Soniat et al., 2013). By close inspection, the xβ-catenin amino acid sequence (665-703) resembles a PY-NLS and is conserved across vertebrate species (Figure 4A). Previous work has demonstrated that the binding affinity of TNPO1 to the PY-NLS is most dependent on the proline (P) and tyrosine (Y) amino acids (or P and methionine (M) in the context of β-catenin) (Cansizoglu, Lee, Zhang, Fontoura, & Chook, 2007). To test the importance of the P and M amino acids for the function of the xβ-catenin NLS, we mutated the codons to two tandem alanine (A) residues and tested how these changes impacted the ability of the xβ-catenin-(665-703) to import a GFP fusion to 3 maltose binding proteins. MBP(x3)-GFP is constitutively excluded from the nucleus due to its large molecular weight (149 kDa) (Figure 4C) (Popken et al., 2015). Fusion of the xβ-catenin-NLS can confer nuclear localization of this large fusion protein. Further, this localization is dependent on the PM motif as substitution of PM with AA abolishes nuclear localization (Figure 4C). These amino acids were also critical for human (h)β-catenin-(665-782)-GFP nuclear accumulation in human cell lines as the PM to AA substitution reduced the mean N:C ratios from 2.5 to 1.5 (Figure 4D). Thus, the PM sequence in the β-catenin NLS is required for nuclear import in yeast and human cells.

**Figure 4.**
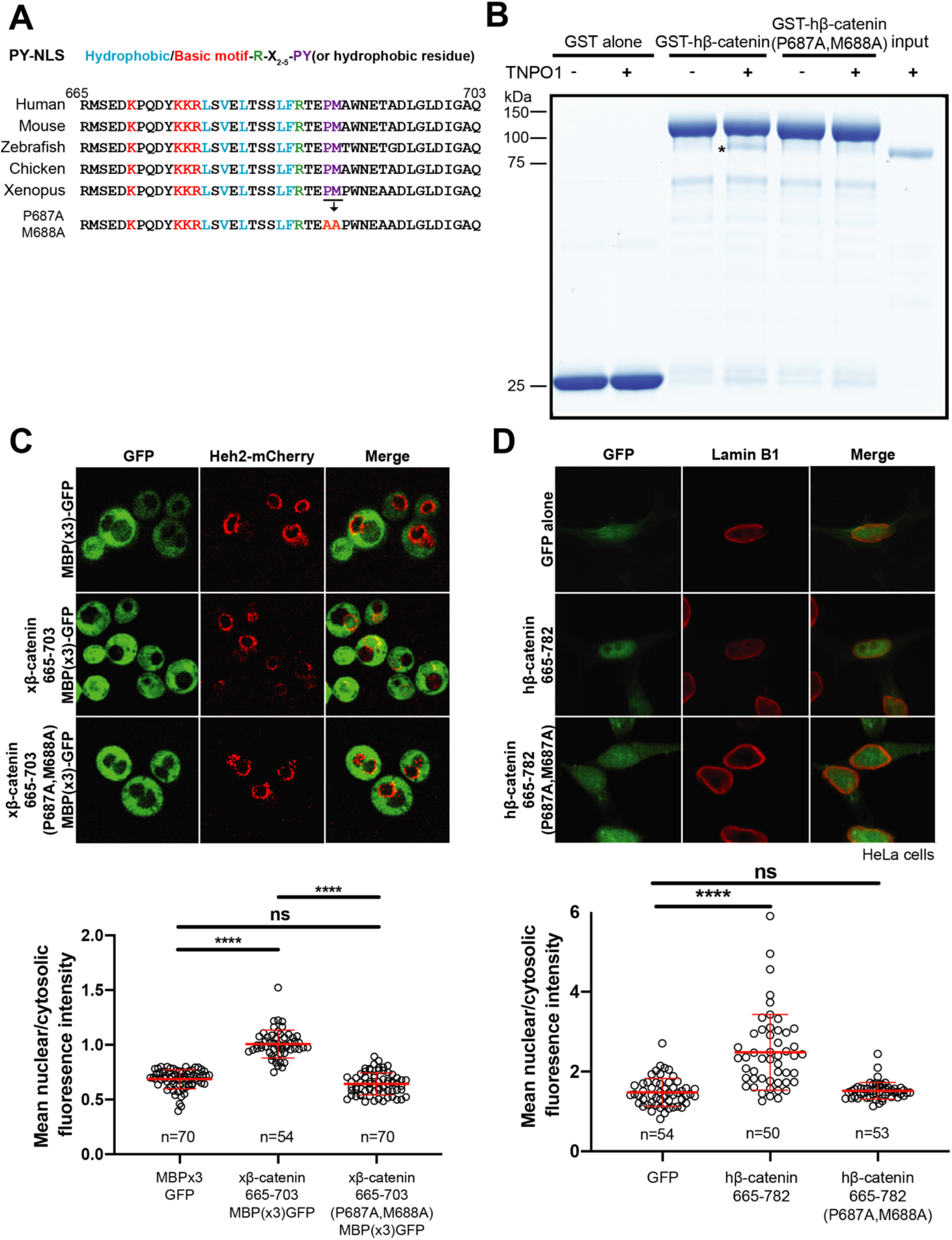
β-catenin contains a PY-like NLS that is required for direct binding to TNPO1. **(A)** Conservation of amino acid sequences that resembles a PY-NLS in the C-terminus of β-catenin. **(B)** In vitro binding assay of purified recombinant TNPO1 to GST fusions of human β-catenin and human β-catenin containing the PM to AA mutations. Proteins were separated by SDS-PAGE and stained with Coomassie Blue. GST alone was used as a negative control. * indicates TNPO1 bound to GST-hβ-catenin. **(C)** Deconvolved fluorescence image of yeast cells expressing MBP(x3)-GFP tagged with the Xenopus β-catenin NLS (665-703) and also an NLS that contains the PM to AA mutation (top). Untagged MBP(x3)-GFP was used as a control. Plot of the ratio of mean nuclear to cytoplasmic fluorescence intensity from a single experiment (bottom) **(D)** Representative fluorescence image of HeLa cells expressing human β-catenin NLS (665-782) or the PM to AA mutant version (top). LaminB1 was labeled to locate the nuclear membrane. GFP alone was used as a control. Plot of the ratio of mean nuclear to cytoplasmic fluorescence intensity from three experiments (bottom). p-values are from unpaired two-tailed t-test where ns is p>0.05, and ****p<0.0001 for both **(C)** and **(D)**.

To confirm that the β-catenin-NLS is directly recognized by TNPO1, we generated recombinant TNPO1 and GST fusions of human β-catenin (GST-hβ-catenin) and human β-catenin containing the PM-AA mutations (GST-hβ-catenin P687A,M688A). We immobilized these GST fusions (and GST alone) on GT Sepharose beads and tested binding to purified TNPO1. We observed specific binding of TNPO1 to the GST-hβ-catenin (Figure 4B), which was disrupted by the PM to AA mutations in the NLS. When taken together, these data establish a model in which TNPO1 can import β-catenin through a direct interaction with its PY-like NLS.

### TNPO1/2 and the β-catenin NLS are required for Wnt signaling *in vivo*

To explore the function of TNPO1-mediated import of β-catenin in vertebrates, we applied two different Wnt signaling assays: 1) a TCF/LEF reporter and 2) *Xenopus* secondary axis development. First, once β-catenin enters the nucleus, it binds to the TCF/LEF complex to activate transcription of Wnt responsive genes (Molenaar et al., 1996; van de Wetering et al., 1997; van de Wetering, Oosterwegel, Dooijes, & Clevers, 1991). A well-established reporter assay (commonly known as TOPFLASH) places the TCF/LEF DNA binding element upstream of a reporter such as GFP or luciferase (Molenaar et al., 1996; van de Wetering et al., 1991). In *X. tropicalis*, the *Tg(pbin7Lef-dGFP)* line has seven tandem TCF/LEF DNA binding sites upstream of GFP and is an effective reporter of Wnt signaling(Borday et al., 2018; Denayer, Tran, & Vleminckx, 2008). We crossed heterozygous transgenic animals with a wildtype animal such that half of the resultant progeny had the transgene. In vertebrates, the *tnpo1* gene is duplicated (*tnpo1* and *tnpo2*), and both paralogs have nearly identical sequence and function (Figure 5-figure supplement 1) (Dormann et al., 2010; Rebane, Aab, & Steitz, 2004; Twyffels, Gueydan, & Kruys, 2014). Therefore, we injected sgRNAs targeting both *tnpo1* and *tnpo2* with Cas9 protein at the one cell stage and raised embryos to st10 before fixing them. Because GFP fluorescence is undetectable at these early stages, we used whole mount *in situ* hybridization (WMISH) to visualize GFP transcripts as an assay for Wnt reporter activation and used sibling embryos without the transgene as a WMISH negative control. When we depleted both *tnpo1* and *tnpo2* using F0 CRISPR, significantly more embryos had weak expression of the GFP transgene compared to uninjected control embryos (Figure 5A, see images of embryos stained for GFP transcripts as key to histogram). This result was specific as a second set of non-overlapping sgRNAs gave comparable results (Figure 5A, *tnpo1*/2 sgRNAs#2). Importantly, we detected deleterious gene modification at the appropriate targeted sites using Inference of CRISPR Edits (ICE) analysis (Figure 5-figure supplement 2).

**Figure 5.**
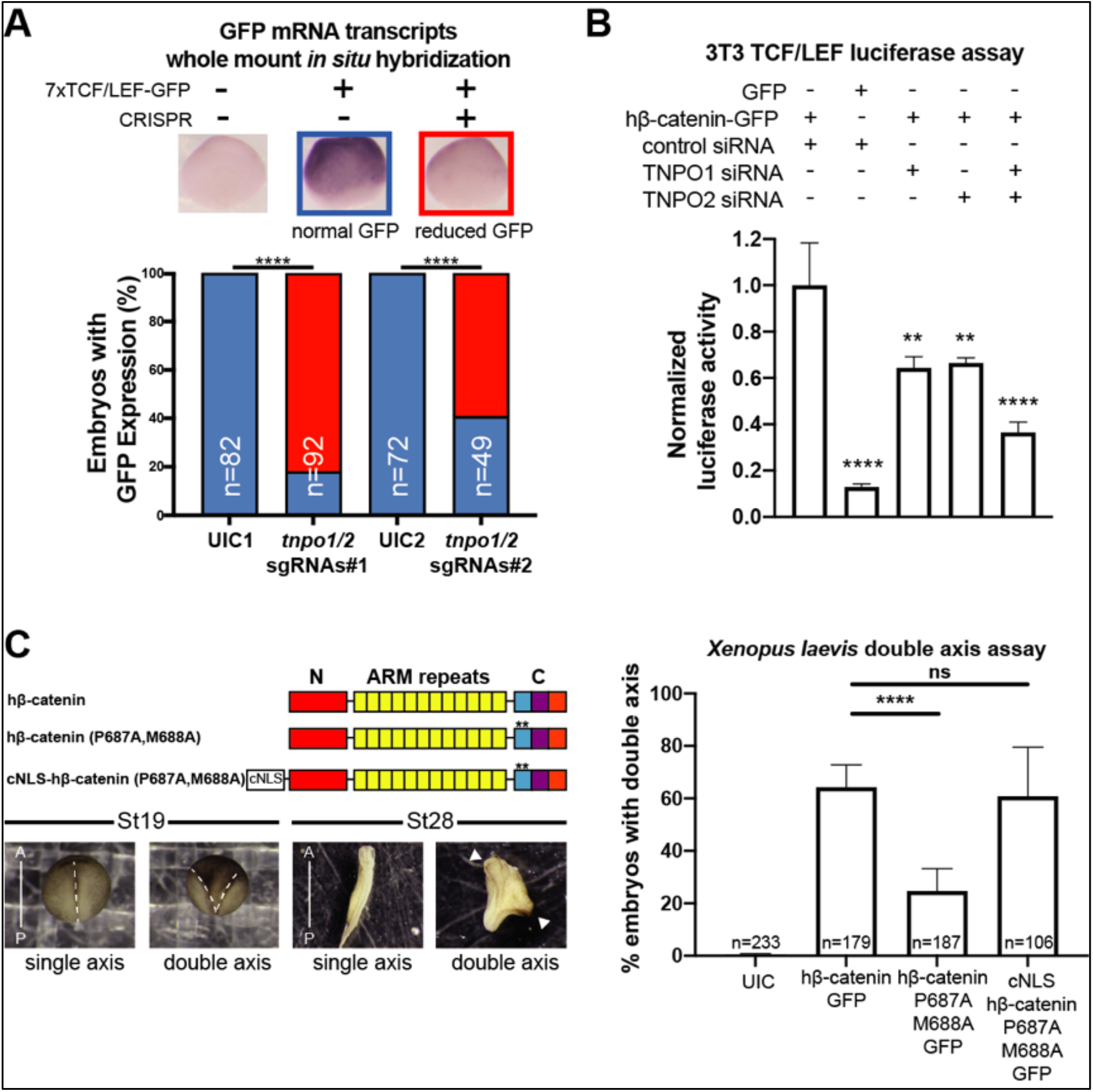
TNPO1/2 and the β-catenin NLS are required for Wnt signaling *in vivo*. **(A)** Depletion of tnpo1 and tnpo2 using two different pairs of non-overlapping sgRNAs represses gfp expression in X. tropicalis Tg(pbin7Lef-dGFP) embryos at stage 10. Key used to quantify embryos with WMISH signal (blue – normal gfp signal, red – reduced gfp signal). Uninjected (UIC) embryos were used as a negative control. **(B)** siRNA mediated TNPO1 and/or TNPO2 knockdown reduces luciferase activity in mouse embryonic fibroblasts that harbor a stable integration of luciferase under the control of TCF/LEF promoters. Wnt signaling was activated by human β-catenin-GFP overexpression. Control siRNA and GFP were used as negative controls. Experiments were performed in triplicate. **(C)** Schematic diagram of three β-catenin constructs used in the double axis assay in Xenopus laevis. ** indicates P687A, M688A substitutions (top left). Dorsal views of X. laevis embryos with anterior to the top (bottom left). Dotted lines indicate the embryonic axis and the white arrows indicate the head. Histogram of the percent of embryos with secondary axes from three independent replicates. (p-values are from Fisher’s exact test **(A)** and **(C)** and unpaired two-tailed t-test **(B)** where ns is p>0.05, p<0.05 (*), 0.0021 (**), 0,0002 (***) and p<0.0001 (****).

We next tested the function of mouse Tnpo1/2 in a stable transgenic mouse fibroblast cell line in which luciferase is expressed under the control of TCF/LEF DNA binding elements. To activate Wnt signaling, we transfected a full length human β-catenin-GFP that increased luciferase signal 7.7 fold over transfection of GFP alone (Figure 5B). Then we measured luciferase activity in transgenic fibroblasts transfected with hβ-catenin-GFP in which we depleted transcripts of TNPO1 or TNPO2 (alone or simultaneously) using specific siRNAs. Compared to control siRNA, depletion of either TNPO1 or TNPO2 led to a 34% reduction in luciferase signal (Figure 5B). By targeting both transcripts, the luciferase signals was reduced by 64% (Figure 5B). By Western blot, we observed the production of hβ-catenin-GFP and the specific reduction of TNPO1/2 (Figure 5-figure supplement 3). Thus, TNPO1/2 are required for robust Wnt signaling.

As reduction of TNPO1/2 would affect the localization of all of their cargos, we next evaluated the specific impact of inhibiting β-catenin nuclear import by testing the function of the PY-AA mutant in the *Xenopus* secondary axis assay. We compared the number of secondary axes induced by the wildtype hβ-catenin mRNA to a PM to AA mutated version (P687A, M688A). We noted a significant reduction in the number of secondary axes induced by the PM-AA mutant β-catenin (Figure 5C). If we add the cNLS to the N-terminus of the PM-AA mutant, then induction of secondary axes is rescued suggesting that the PM-AA mutant fails to enter the nucleus to activate Wnt signaling (Figure 5C).

### The M9M peptide inhibits Wnt signaling

Having established that TNPO1 binds directly to a PY-like NLS and imports β-catenin into the nucleus, we wondered whether direct perturbation of the β-catenin-TNPO1 interaction could be a viable therapeutic strategy. We therefore took advantage of the prior design of a potent TNPO1 peptide inhibitor, M9M, that binds with high affinity to the TNPO1 NLS binding site (Cansizoglu et al., 2007). We tested whether this peptide could inhibit Wnt signaling in the mouse fibroblast TCF/LEF luciferase reporter cell line (Cansizoglu et al., 2007). Excitingly, transfection of the M9M peptide reduced luciferase activity in a dose dependent manner, regardless of whether activation was induced with a Wnt ligand (Wnt3a) or by co-transfection with human β-catenin (Figure 6A, Figure 6-figure supplement 1A-B).

**Figure 6.**
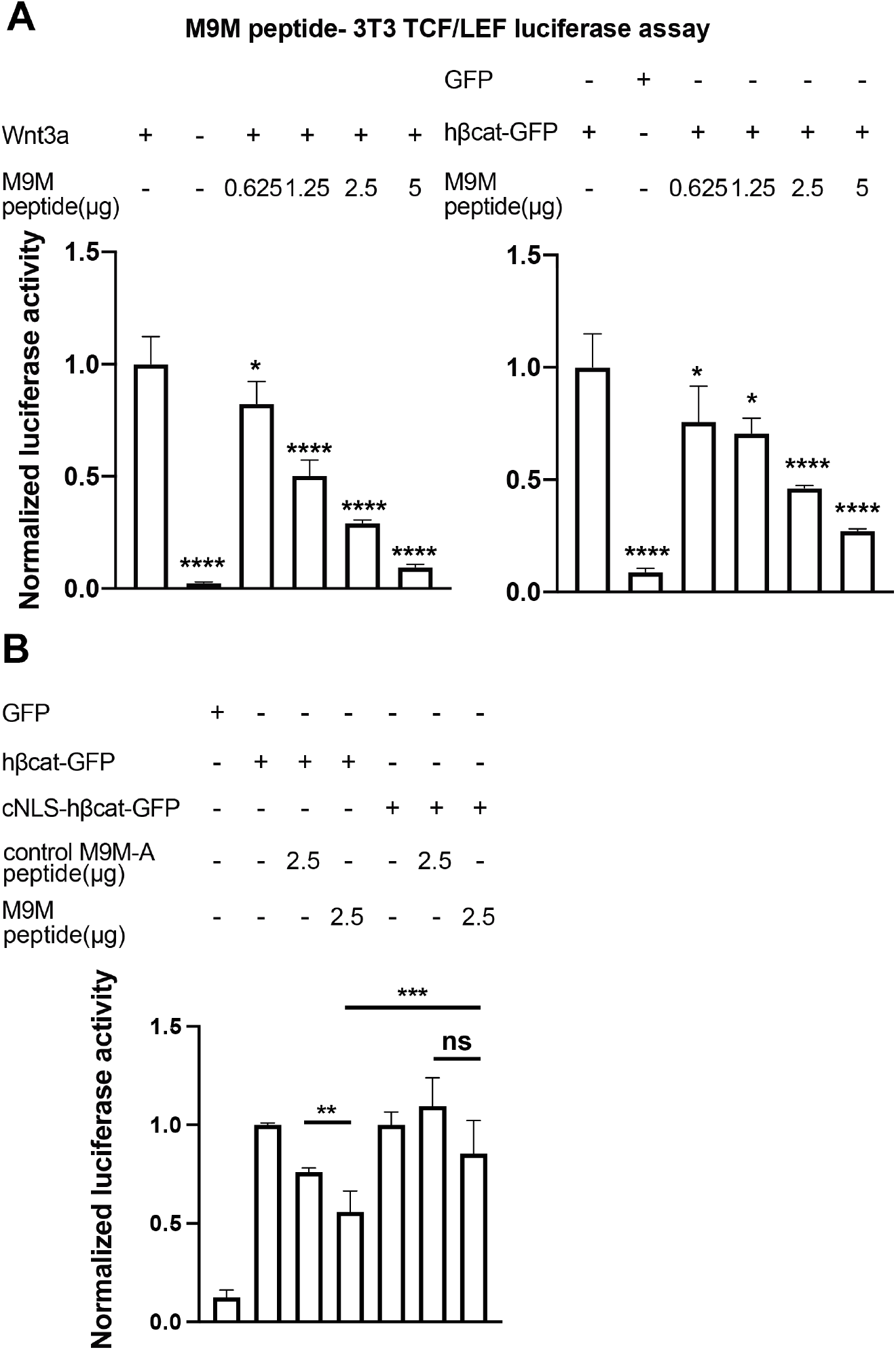
The M9M peptide inhibits Wnt signaling. Wnt signaling was activated by Wnt3a (A, left), human β-catenin-GFP overexpression (A, Right and B) or cNLS-human β-catenin-GFP (B). No Wnt3a or GFP overexpression were used as negative controls. Experiments were performed in triplicate (A) or duplicate in two independent experiments (B). (p-values are from unpaired two-tailed t-test where ns is p>0.05, p<0.05 (*), 0.0021 (**), 0.0002 (***) and p<0.0001 (****).

The M9M peptide is a chimera of the NLSs of hnRNP M and A1. To test specificity, we leveraged the understanding of the key amino acids that confer binding to TNPO1 in each individual NLS to design a control M9M-A peptide. M9M-A contains 7 amino acid substitutions that would abrogate binding to TNPO1 (Figure 6-figure supplement 1C). The M9M-A peptide only reduced luciferase activity by 24% (compared to 45% by M9M at the similar dosage; Figure 6B).

As the M9M peptide inhibits the nuclear import of a multitude of TNPO1/2 cargos (Cansizoglu et al., 2007), we sought to ensure that the M9M-mediated inhibition of Wnt signaling was specifically due to the reduced nuclear import of β-catenin. We therefore transfected a human β-catenin with a cNLS, which would be imported by Kap-α/Kap-β1 and thus be insensitive to M9M inhibition. The cNLS-human β-catenin could drive the luciferase reporter to levels comparable to human β-catenin but the M9M peptide only reduced this signal by ∼15% (Figure 6B). Thus, these data support the conclusion that the M9M peptide’s impact on Wnt signaling is due, at least in part, to the inhibition of β-catenin nuclear transport via TNPO1/2. Together, Wnt signaling can be inhibited by blocking TNPO1 mediated import of β-catenin either by mutating the β-catenin NLS or competitive inhibition with the M9M peptide.

## DISCUSSION

Our lack of understanding of the mechanism of β-catenin nuclear transport has been a major stumbling block for opening up new avenues of Wnt-targeted anti-cancer therapies. Numerous transport models have been proposed including piggybacking on TCF/LEF (Behrens et al., 1996; O. Huber et al., 1996; Molenaar et al., 1996) or APC (Henderson, 2000; Neufeld, Zhang, Cullen, & White, 2000; Rosin-Arbesfeld, Townsley, & Bienz, 2000) although neither factor was ultimately found to be required for import (Eleftheriou, Yoshida, & Henderson, 2001; Orsulic & Peifer, 1996; Prieve & Waterman, 1999). There is also a model in which β-catenin directly binds to the FG-nups and translocates through the NPC in a NTR-like (yet Ran independent) mechanism (Fagotto, 2013; Xu & Massague, 2004). Indeed, while some of our data might hint that there are elements in β-catenin that might have an affinity for the NPC (Figure 2D, white arrows), by far the major determinant of its nuclear accumulation is the PY-like NLS in its C-terminus. Thus, the most parsimonious mechanism of β-catenin nuclear import is also the most straightforward - it is a Kap-β2/TNPO1 cargo. This conclusion is based on 1) The identification of a PY-like NLS in the C-terminus of β-catenin, 2) The direct binding between TNPO1 and this NLS *in vitro* and 3) The essential role for this NLS sequence and TNPO1/2 in mediating TCF/LEF binding activation and secondary axis formation of β-catenin in multiple model systems.

This work must also be reconciled with a recent study that implicated Importin 11 as an important factor for β-catenin nuclear transport in a subset of cancer cells (Mis et al., 2020). The latter work also identified the C-terminus as essential for β-catenin nuclear transport, and our study adds further resolution to a specific PY-like NLS. However, we did not determine any requirement for Importin 11 (Kap120) in β-catenin import, at least in the yeast system. Although it remains possible that Importin 11 might ultimately have a role in β-catenin nuclear transport in some vertebrate cells, the current evidence suggests that it plays a critical role only in a subset of colon cancer cells (Mis et al., 2020). In contrast, our work establishes that TNPO1 imports β-catenin in yeast, amphibian, mouse and human cells, providing confidence that there is a TNPO1-specific mechanism at play. How importin 11 specifically contributes to β-catenin nuclear import in the context of specific cancer cell lines remains to be fully established.

For the purposes of rational drug design, we fortuitously identified the β-catenin NTR as TNPO1, as it is one of the few NTRs where the NLS-NTR interaction is resolved to the atomic level (Cansizoglu et al., 2007; Soniat et al., 2013). This knowledgebase has established a consensus amino acid sequence (PY-NLS) that helped us identify the β-catenin NLS and the key PM residues required for TNPO1 binding. Further, it has led to the generation of the high affinity M9M peptide (Cansizoglu et al., 2007) that, as shown here, can be used to inhibit Wnt signaling in TCF/LEF luciferase mouse fibroblast cell lines, demonstrating proof of principle that inhibiting TNPO1 could be a viable therapeutic strategy. Nonetheless, such a strategy might benefit from future crystallographic studies of the TNPO1-β-catenin complex as the overall binding of different NLSs can vary depending on the contribution of individual amino acids. It may be possible, for example, to identify small molecules that specifically block the β-catenin-TNPO1 interaction without more broadly impacting other TNPO1 cargos. Our findings will therefore form the basis for future work to identify small molecules that could specifically target and ameliorate the multitude of Wnt related diseases including cancers.

## ACKNOWLEDGEMENTS

We thank M. Slocum and M. Lane for *Xenopus* husbandry and E. Rodriguez, N. Ader, and S. Chandra for technical assistance and advice. For providing plasmids, we thank Dr. Yuh Min Chook at UTSW and Addgene. We thank the National Xenopus Resource at the Marine Biological Laboratory for distributing the *X. tropicalis Tg(pbin7Lef-dGFP)* line. We thank the Yale Center for Advanced Light Microscopy for their assistance with confocal imaging. W.Y.H was supported by the Paul and Daisy Soros Fellowship for New Americans and the NIH (5F30HL143878). W.Y.H., and V.K were supported by the Yale MSTP NIH T32GM07205. D.P.G was supported by the NIH (5F31HL149246). M.K.K. and C.P.L were supported by the NIH (2R01HL124402).

## CONTRIBUTIONS

W.Y.H., C.P.L., and M.K.K. designed the study and wrote the manuscript which was read and approved by all authors. W.Y.H., and V.K performed the *Xenopus* experiments. W.Y.H., and D.P.G performed yeast experiments. W.Y.H performed mammalian cell experiments.

## DECLARATION OF INTERESTS

M.K.K is a co-founder of Victory Genomics, Inc.

## MATERIALS AND METHODS

### CONTACT FOR REAGENT AND RESOURCE SHARING

Further information and request for reagents may be directed to and will be fulfilled by the Lead Contact, Mustafa K. Khokha

### KEY RESOURCES TABLE

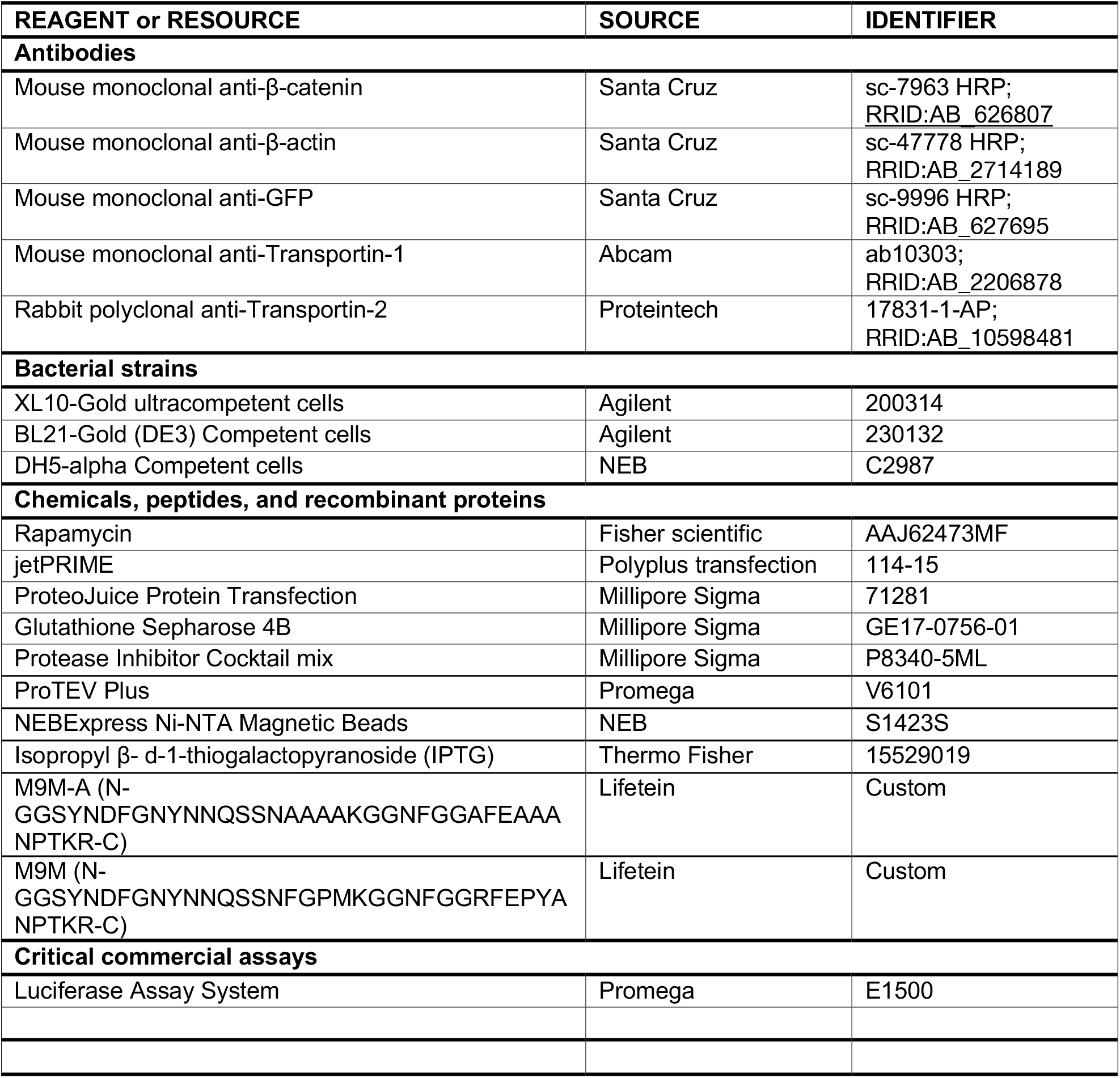
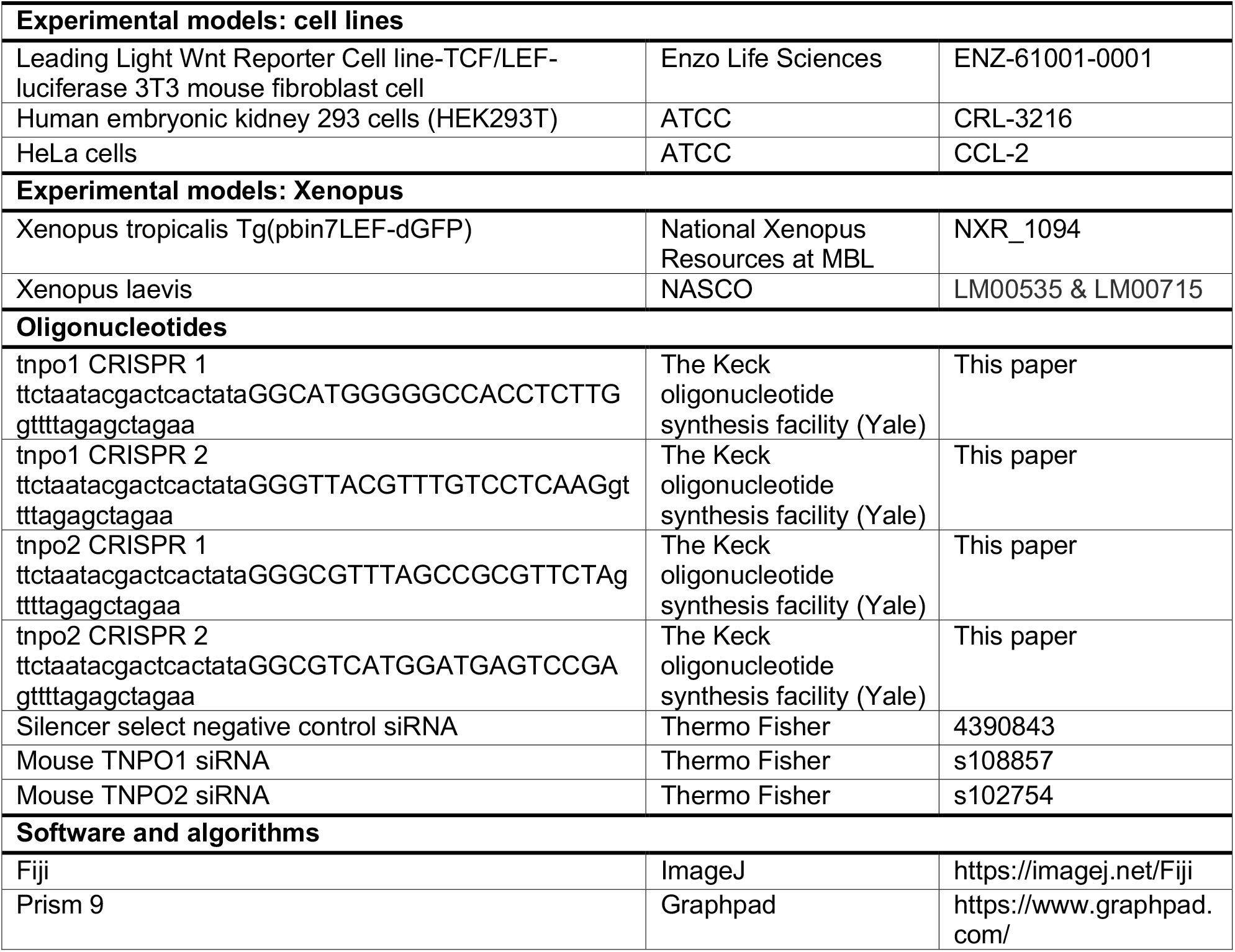

**Table S1.**
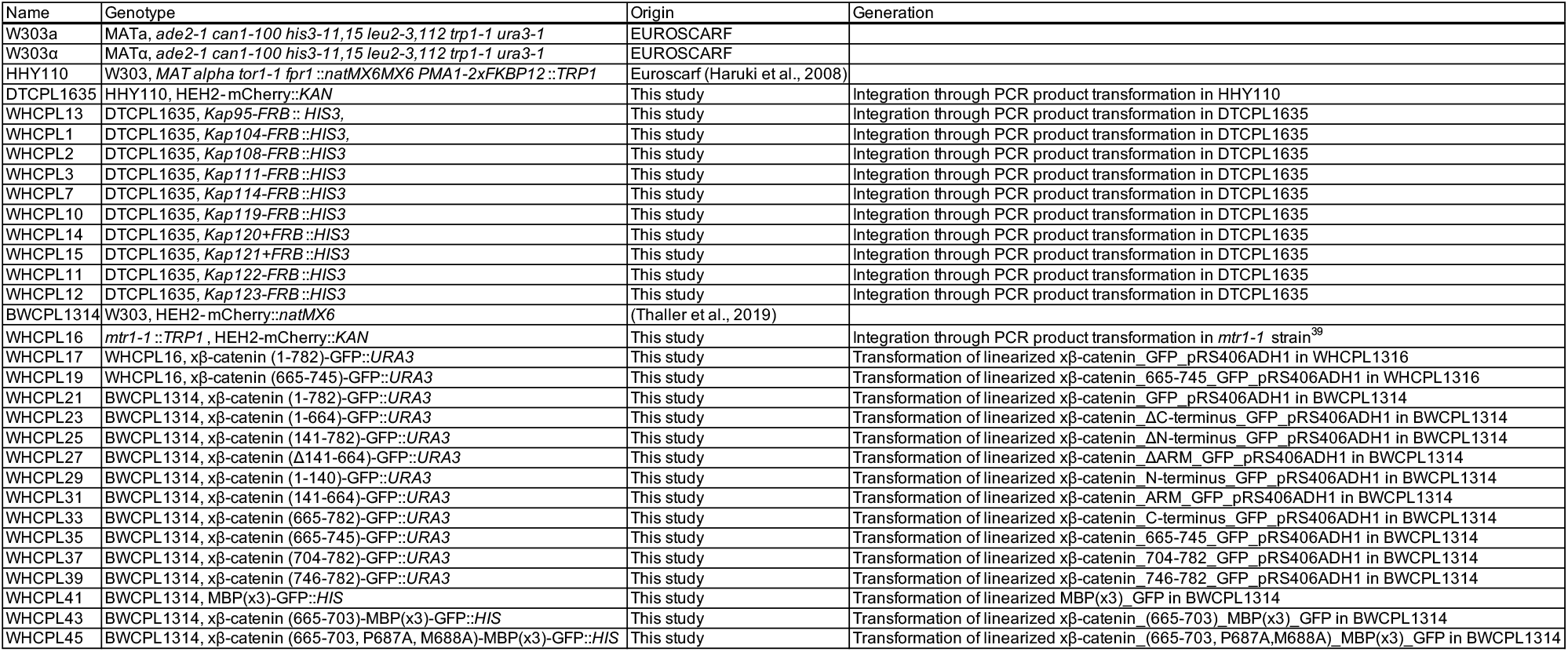
*S. cerevisiae* strains.

**Table S2.**
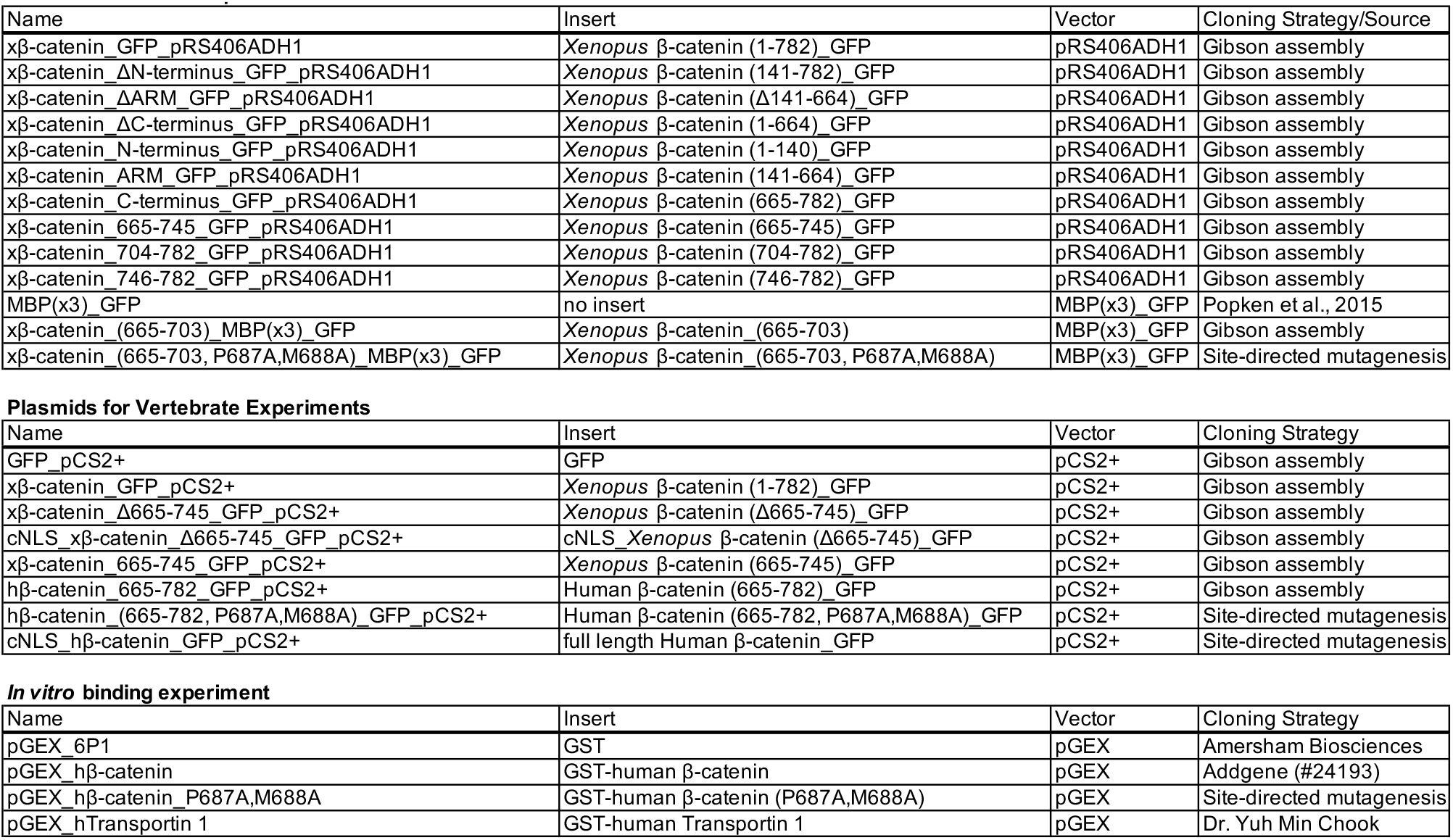
Plasmids.

**Table S3.**
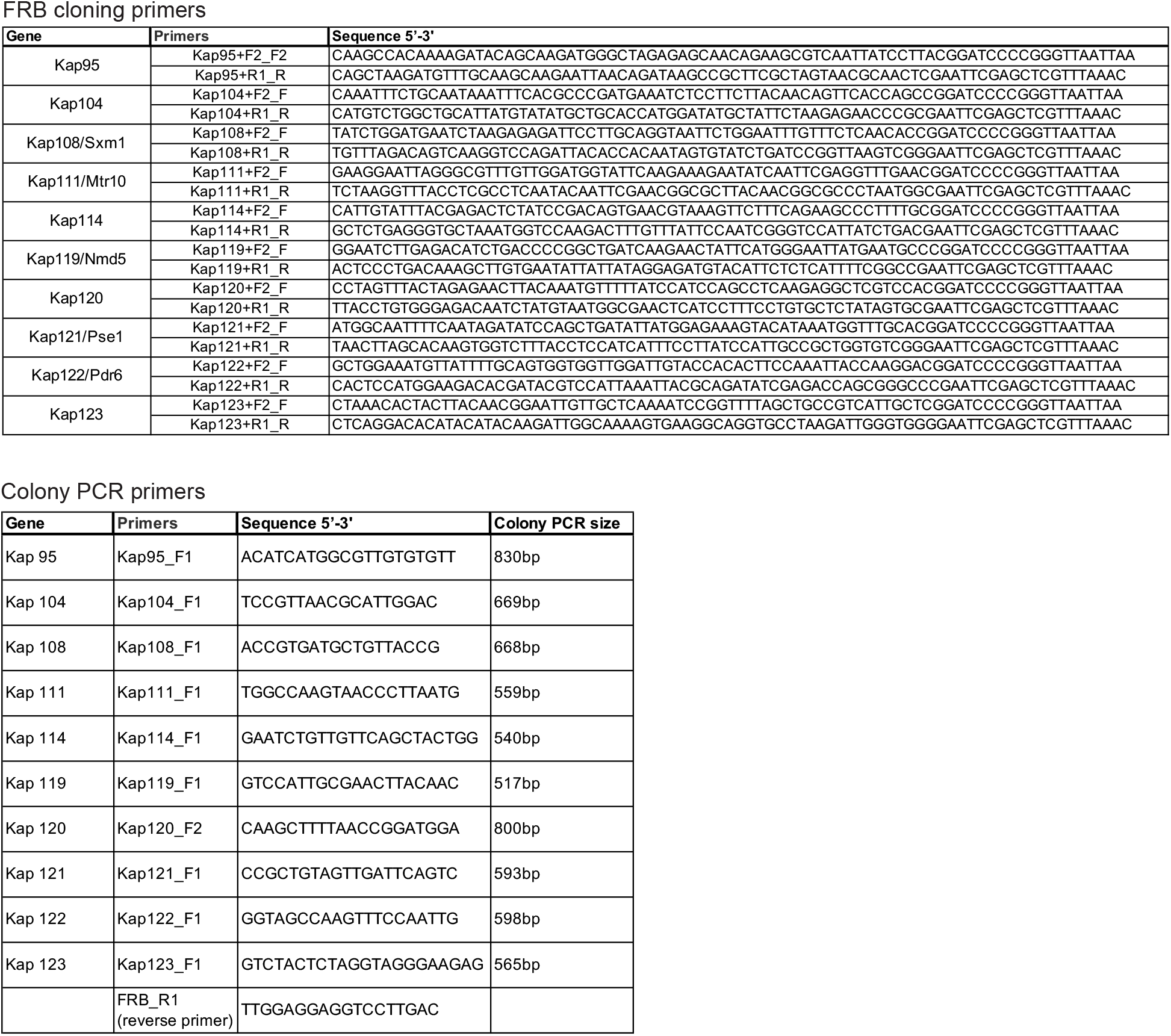
Anchor away primers.

### EXPERIMENTAL MODEL AND SUBJECT DETAILS

#### Xenopus

*Xenopus tropicalis* and *Xenopus laevis* were maintained and cared for in accordance to the Yale University Institutional Animal Care and Use Committee (IACUC) protocols. *In vitro* fertilization was performed as per standard protocols (del Viso & Khokha, 2012; Sive, Grainger, & Harland, 2007).

#### *Saccharomyces cerevisiae* strains

All yeast strains used in this study are listed in Table S1. Yeast strains were grown at 30°C, unless indicated otherwise in YPD medium (1% Bacto yeast extract, 2% bactopeptone, 2% glucose, 0.05% Adenine sulfate). Transformation of yeast was carried out using standard protocols (Amberg, 2005).

#### Mammalian cells

HEK293T and HeLa cells were maintained and cultured with DMEM medium + 10% Fetal Bovine Serum (FBS) + 1% Penicillin and Streptomycin (PS) in a T-75 flask. The engineered 3T3 mouse fibroblast cell line that stably expresses the TCF/LEF luciferase transgene was maintained with DMEM medium + Cell Growth Medium Concentrate (Enzo life sciences). Further experimental procedures for the 3T3 cell line can be found under TCF/LEF luciferase assay method section. Upon 70-80% cell confluency, cells were transfected with plasmids using jetPRIME (Polyplus-transfection) following the manufacturer’s instructions. Cells were fixed with 4% paraformaldehyde/PBS and further processed for immunofluorescence 24-48 hours post-transfection. Antibodies used in this study are listed in Key Resources Table.

### Method details

#### Plasmid, mRNA, siRNA, CRISPR and M9M peptide

Key resources and all plasmids used in this study are listed in Key Resources Table and Table S2, respectively. *Xenopus* β-catenin-GFP (Addgene #16839), *Xenopus* cNLS-β-catenin-GFP (Addgene #16838), GST-human β-catenin (Addgene #24193), and NLS-mCherry (Addgene #49313) plasmids were obtained from Addgene. GST-transportin 1 (TNPO1) and Gal-MBP(x3)-GFP plasmids were generous gifts from Dr. Yuh-min Chook at UTSW and Dr. Liesbeth M. Veenhoff at University of Groningen, respectively. Gibson Assembly (New England Biolabs) was used to generate GFP tagged *Xenopus* β-catenin truncation constructs following the manufacturer’s instructions. Both the human and *Xenopus* β-catenin P687A, M688A variants were generated using Q5 site-directed mutagenesis (New England Biolabs) following the manufacturer’s instructions. Subsequently, the β-catenin constructs were sub-cloned into the pRS406 vector containing an ADH1 promotor for yeast studies or a pCS2+ vector for mammalian/*Xenopus* studies. mRNAs were generated using the SP6 mMessage machine kit (Thermo Fisher Scientific) and RNA clean & concentrator kit (Zymo Research) following the manufacturer’s instructions. We obtained siRNAs directed against mouse TNPO1 (s108857), mouse TNPO2 (s102754) and a control siRNA (4390843) from Thermo Fisher Scientific. To generate CRISPR sgRNAs, we used the EnGen sgRNA synthesis kit (NEB) following the manufacturer’s instructions with the following targeting sequences *tnpo1* sgRNA#1 (5’-GGCATGGGGGCCACCTCTTG-3’), *tnpo1* sgRNA#2 (5’-GGGTTACGTTTGTCCTCAAG-3’), *tnpo2* sgRNA #1 (5’-GGGCGTTTAGCCGCGTTCTA-3’), and *tnpo2* sgRNA #2 (5’-GGCGTCATGGATGAGTCCGA-3’) (designed using CRISPRscan(Moreno-Mateos et al., 2015)). CRISPR experiments in wildtype or transgenic *Xenopus tropicalis* were performed as previously described(Bhattacharya, Marfo, Li, Lane, & Khokha, 2015).CRISPR gene editing efficiency was assessed using Synthego ICE (ice.synthego.com) as previously described(Sempou, Lakhani, Amalraj, & Khokha, 2018). The M9M peptide (GGSYNDFGNYNNQSSNFGPMKGGNFGGRFEPYANPTKR) and the M9M-A peptide (GGSYNDFGNYNNQSSNAAAAKGGNFGGAFEAAANPTKR) were synthesized by LifeTein.

#### Yeast and mammalian β-catenin sub-cellular localization by microscopy

Expression plasmids containing full length and fragments of the β-catenin-GFP coding sequence under the control of the *ADH1* promoter were transformed into the W303, Heh2-mCherry::NAT (BWCPL1314) strain. A yeast colony that incorporated the plasmid sequence was cultured and mounted onto a coverslip for live imaging on a DeltaVision wide-field microscope (GE Healthcare) with a CoolSnapHQ^2^ CCD camera. Yeast fluorescent images were deconvolved using the iterative algorithm sofWoRx. 6.5.1 (Applied Precision, GE Healthcare). β-catenin-GFP transfected HeLa or HEK293T cells were mounted on Pro-Long Gold coated coverslip for imaging on a Zeiss Axio Observer microscope. All fluorescent images were analyzed with Fiji software. For quantification, the oval selection tool was used to draw a circle in both the nuclear and cytosolic regions on the same image plane to measure florescence intensity.

#### Secondary axis assays and β-catenin sub-cellular localization

*Xenopus laevis* embryos were injected with a mixture of either 200pg of *Xenopus* or human β-catenin-GFP mRNA and cNLS-mCherry mRNA in one of four cells (targeting the ventral side). Embryos were assessed for a secondary axis via stereomicroscopy at stage 17-19. For the β-catenin localization experiments by fluorescence, Stage 10 embryos were fixed in 4% paraformaldehyde/PBS at 4°C overnight on a nutator. Embryos were washed in 1x PBS + 0.1% TritonX-100, and the dorsal blastopore lips were sectioned with a razor blade and mounted on Pro-Long Gold (Invitrogen) coated coverslip before imaging on a Zeiss 710 confocal microscope.

#### The Anchor Away assay

To employ the Anchor Away approach, we used a yeast strain that harbors a FKBP12 fusion of the endogenous plasma membrane H^+^-ATPase (*PMA1* gene), a Heh2-mCherry fusion to mark the nuclear envelope, and a mutated *TOR1* gene (HHY110: *HEH2-mCherry::KAN, PMA1-2xFKBP12, fpr1::NAT tor1-1*). In this strain, individual, endogenous NTRs are tagged with the FRB domain at the C-terminus by homologous recombination of a PCR product that contains an FRB sequence, a 3x HA epitope, and a selective marker, *HIS3*, flanked by a 60 bp homology arm of endogenous NTR coding sequence (Figure S8A) (Haruki et al., 2008; Longtine et al., 1998). Integration of an FRB sequence is confirmed by colony PCR using a gene specific forward and a plasmid specific reverse primer and rapamycin induced cell death for essential NTRs (Figure S8B, S8C Table S3 for colony PCR primers). Subsequently, an expression plasmid containing the coding sequence for xβ-catenin (665-782)-GFP under the control of the *ADH1* promoter was transformed into the Anchor Away line. These lines were treated with 1 mg/ml of rapamycin (5-15 minutes of incubation) or vehicle alone (DMSO) before imaging.

#### TCF/LEF Luciferase Assay

A 3T3 mouse fibroblast cell line that has a stable integration of the luciferase reporter gene under the Wnt responsive TCF/LEF promoters were used for this assay (Enzo life sciences). Cells were maintained with DMEM medium + Cell Growth Medium Concentrate (Enzo life sciences). Prior to the transfection, cells were seeded on a 24 well plate in media containing DMEM + Cell Assay Medium Concentrate (Enzo life sciences). Subsequently, cells were transfected with siRNA (25 pmol) first, then GFP or human β-catenin-GFP DNA (0.5 µg) for 48 hours and 24 hours, respectively using JetPRIME per the manufacturer’s instructions. Luciferase was quantified using the Luciferase Assay System (Promega) and the Promega Glomax luminometer according to the manufacturer’s instructions. For the M9M peptide experiment, cells were transfected with an M9M or M9M-A peptide dose ranging from 0.635 µg to 0.5 µg using ProteoJuice Protein transfection following the manufacturer’s instructions for 20 hours and Wnt signaling was activated either by Wnt3a ligand (50 ng/ml) or human β-catenin-GFP DNA (0.5 µg) for 16-24 hours.

#### *In vivo* TCF/LEF GFP *in situ* hybridization

Heterozygous *Xenopus tropicalis Tg(pbin7Lef-dGFP)* were crossed with wild-type *X. tropicalis*. Fertilized embryos were injected with sgRNAs targeting *tnpo1* and *tnpo2* and Cas9 protein at one-cell stage and collected at stage 10 for *in situ* hybridization as previously described (Khokha et al., 2002). Digoxigenin-labeled anti-sense GFP probe was used to detect GFP transcript expression. Progeny that did not carry the transgene were used as a negative control.

#### Western blotting

3T3 mouse fibroblast cells were lysed in RIPA buffer to harvest protein samples. Protein levels are normalized and immunoblots were carried out in Bolt 4%-12% Bis-Tris plus gels following standard protocols. Antibodies used in this study are listed in Key Resources Table.

#### *In vitro* binding experiment

pGEX-6P1, pGEX-human β-catenin or pGEX-human transportin 1 were transformed into the BL21 *E. coli* strain and cultured in LB with antibiotics to mid-log phase (OD_600_ 0.6-0.8). To induce expression of the recombinant proteins (GST alone, GST-hβ-catenin and GST-hTNPO1), IPTG was added at a final concentration of 1 mM for 3 hours. All cultures were harvested in 50 mL batches and stored at -20°C until further use. Glutathione Sepharose (GT) beads (Millipore Sigma) were washed and equilibrated in lysis buffer (50 mM Tris pH 7.4, 150 mM NaCl, 2 mM MgCl_2_, 10% glycerol, 0.05% NP-40, 1 mM DTT and protease inhibitor cocktail mix (Millipore Sigma)). Bacterial pellets were resuspended with the ice cold lysis buffer, sonicated and spun down at 4°C at 30,000 x g for 20 minutes. The supernatant was collected into a new 50 ml conical tube and incubated with 200 µl of GT bead slurry for 1 hour at 4°C. Subsequently, the GST-GT bead slurry was collected and washed with lysis buffer (excluding the protease cocktail mix). The GST tag was removed from hTNPO1 using proTEV Plus Protease (Promega), and the protease enzyme was further removed from hTNPO1 protein by Ni-NTA Magnetic Beads (NEB) per the manufacturer’s instructions. hNTPO1 protein was incubated with GT beads preloaded with GST fusion protein (GST alone or GST-hβ-catenin) for 1 hour at 4°C. The beads were washed with the lysis buffer and eluted with SDS-PAGE sample buffer. Protein samples were separated by SDS-PAGE and detected with Coomassie (BioRad).

#### Quantification and Statistical Analysis

Statistical significance was defined as p<0.05 (*), 0.002 (**), 0.0002 (***), and 0.0001 (****). The double axis assay and *in situ* data were analyzed by Fisher’s exact tests. Otherwise, unpaired two-tailed Student’s t tests or a one way ANOVA test was used to determine significance of mean ratio of nuclear to cytosolic fluorescence intensity in GraphPad Prism 8.4.3.

---

**Figure 2-figure supplement 1.**
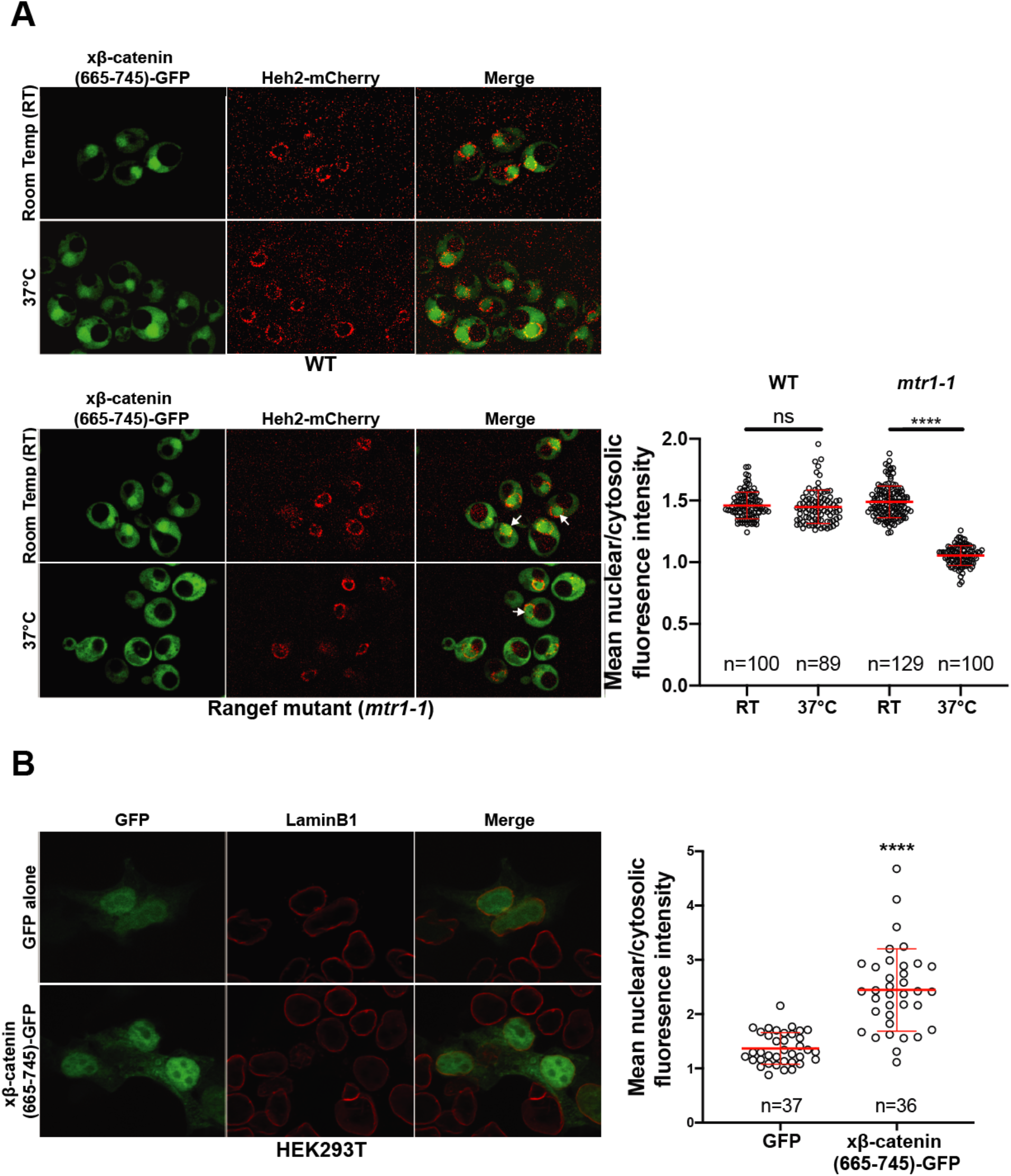
Sub-cellular localization of *Xenopus* β-catenin (665-745)-GFP in the *S. cerevisiae mtr1-1* strain and HEK293T cells. **(A)** Deconvolved fluorescence image of *Xenopus* β-catenin-(665-745)-GFP in the wild-type (top left) and RanGEF mutant (*mtr1-1*) (bottom left) strain at room temperature (RT) or 37°C that co-expresses Heh2-mCherry as a nuclear envelope marker. White arrows indicate the nuclear compartment. The ratio of mean nuclear to cytosolic fluorescence intensity from a single experiment (right) **(B)** Representative image of HEK293T cells expressing *Xenopus* β-catenin (665-745)-GFP. LaminB1 was labeled to locate the nuclear envelope. GFP was used as a control. Ratio of nuclear to cytoplasmic intensities from two independent replicates (right). *p*-values are from unpaired two-tailed t-test where ns is p>0.05, and ****p<0.0001.

**Figure 2-figure supplement 2.**
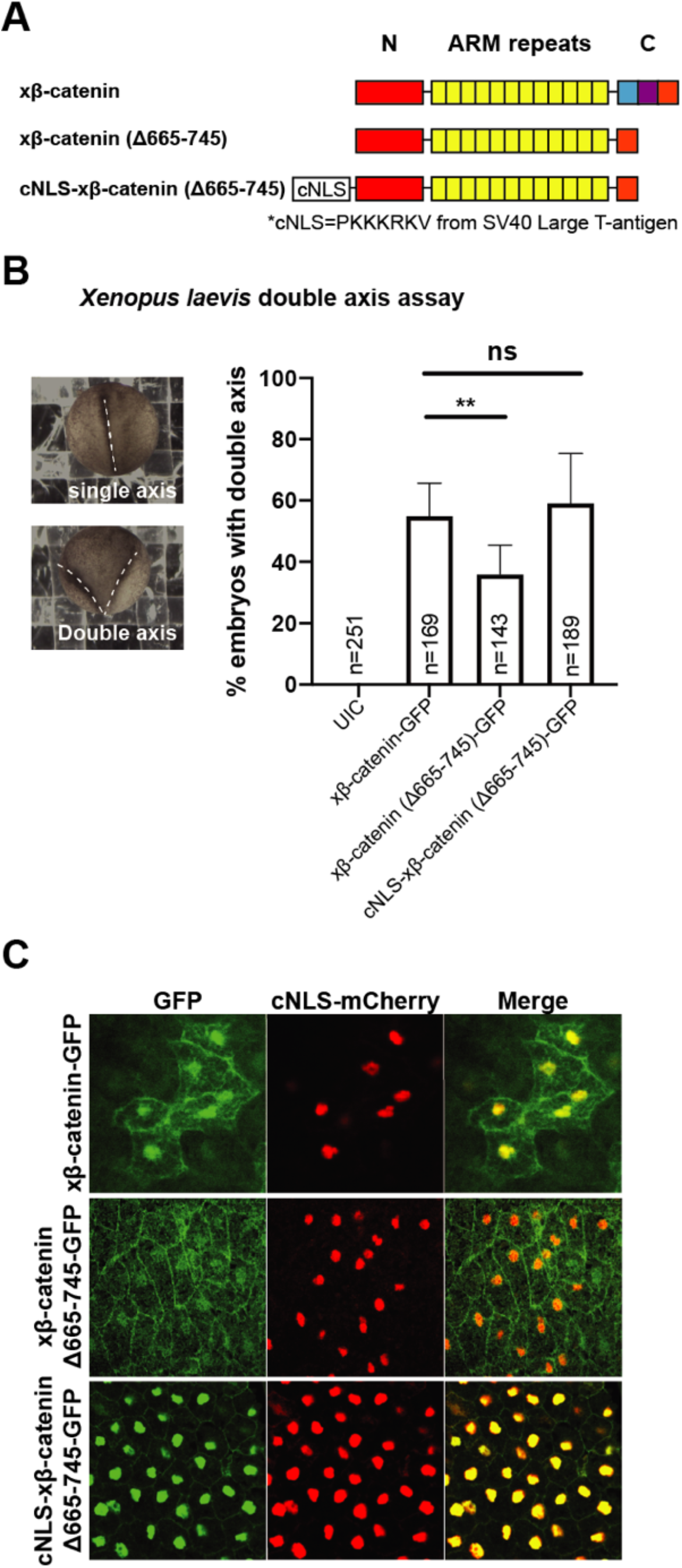
Residues 665-745 of β-catenin are required to induce secondary axes in *Xenopus laevis*. **(A)** Schematic diagram of *Xenopus* β-catenin constructs **(B)** Double axes were scored in st 19 embryos viewed dorsally with anterior to the top (left). Data from three independent replicates depicted in histogram (right). *p*-values are from Fisher’s exact test where ns is p>0.05, p<0.05 (*), and 0.0021 (**). **(C)** Subcellular localization of *x*β-catenin-GFP, *x*β-catenin (Δ665-745)-GFP or cNLS-*x*β-catenin (Δ665-745)-GFP in the dorsal blastopore lip of stage 10 *Xenopus laevis* embryos. cNLS-mCherry mRNA was co-injected to mark the nucleus.

**Figure 3– figure supplement 1.**
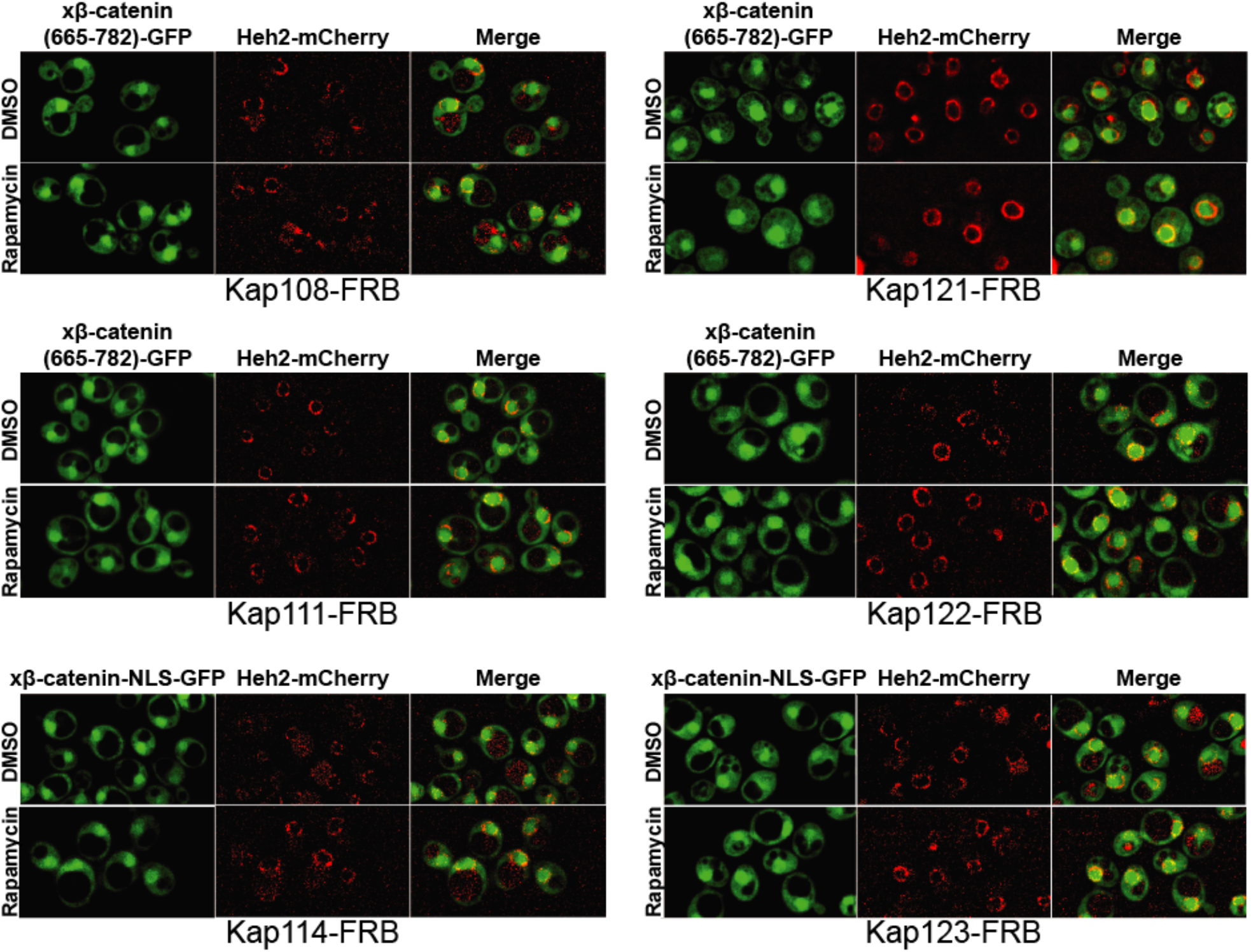
Sub-cellular localization of *Xenopus* β-catenin (665-782)-GFP in Anchor Away strains in *S. cerevisiae*. Representative deconvolved fluorescence image of *x*β-catenin (665-782)-GFP treated with DMSO (carrier) or rapamycin in the indicated NTR-FRB strain. Heh2-mCherry was used as a nuclear membrane marker.

**Figure 5-figure supplement 1.**
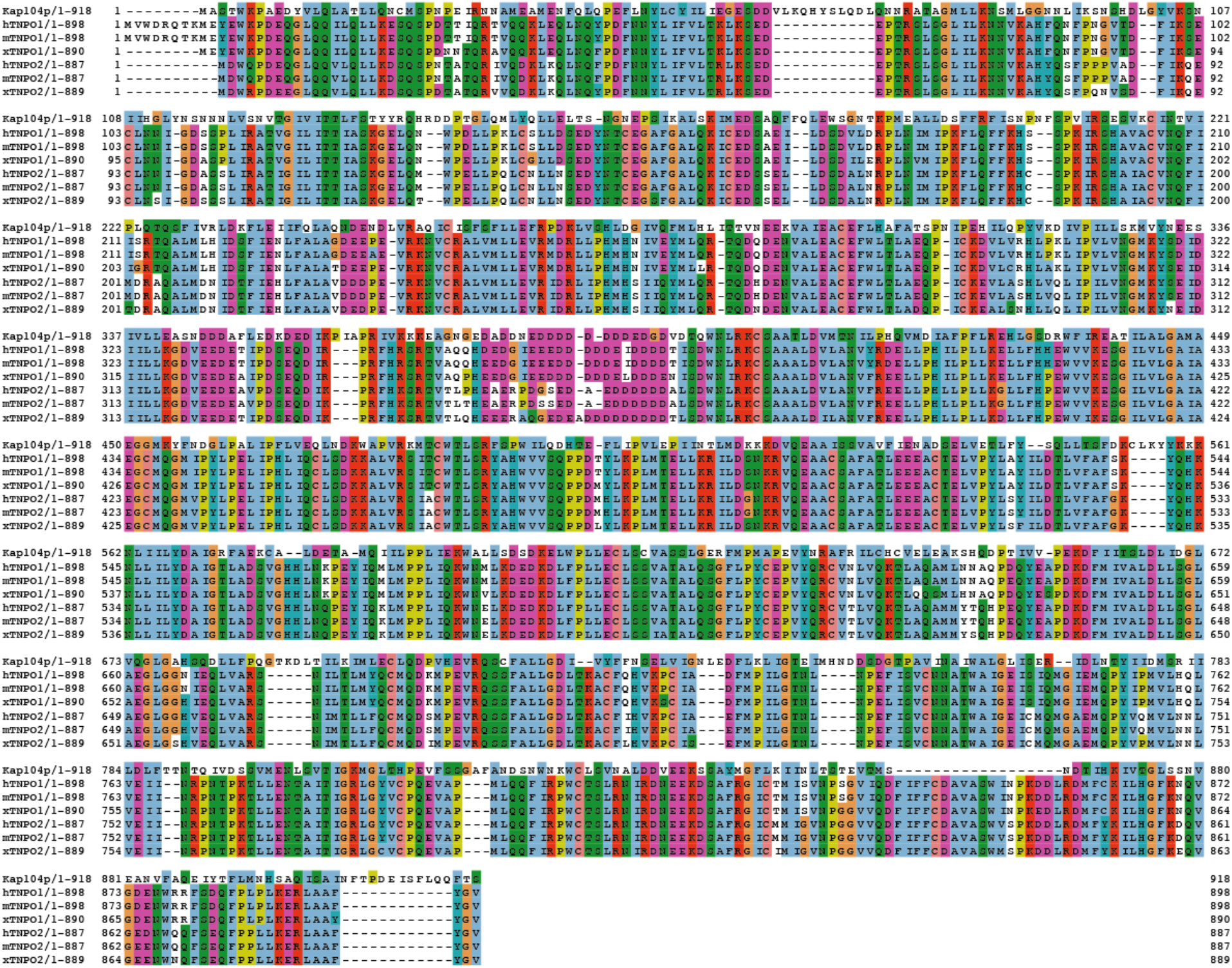
Sequence alignment of Tnpo1 and Tnpo2 across species. Amino acid sequences of transportin 1 and transportin 2 were compared across four different species (*S. cerevisiae*, human, mouse and *Xenopus tropicalis)*. Each residue in the alignment is colored using Jalview software(Waterhouse, Procter, Martin, Clamp, & Barton, 2009).

**Figure 5-figure supplement 2.**
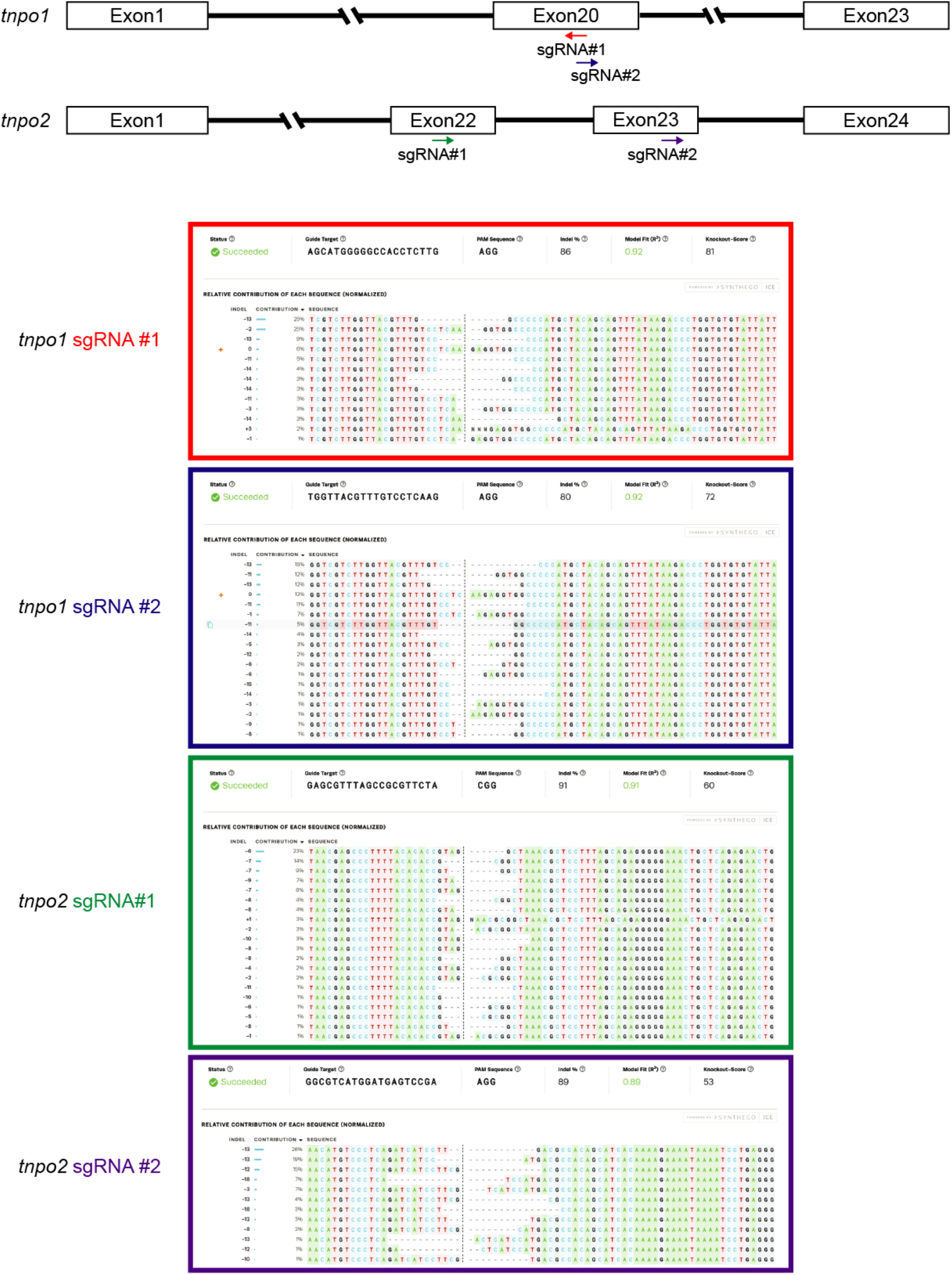
*X. tropicalis tnpo1* and *tnpo2* gene depletion by CRISPR/Cas9. Schematic diagram of *tnpo1* and *tnpo2* sgRNA target sites (top). ICE analysis of indel mutations at predicted target sites (bottom).

**Figure 5-figure supplement 3.**
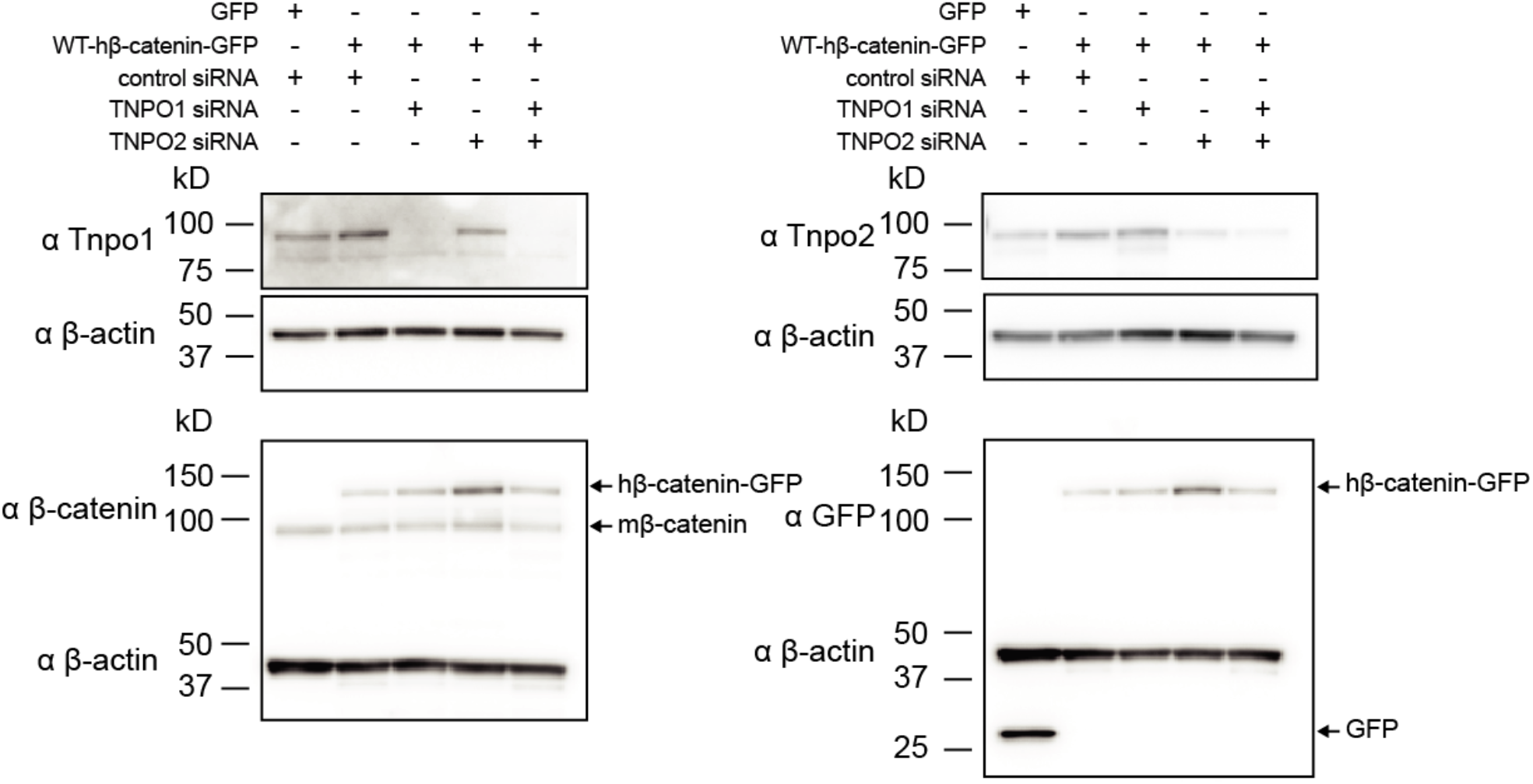
Western blots of Tnpo1/2 and β-catenin from 3T3 TCF/LEF luciferase assays. Western blot demonstrating the efficacy of siRNA mediated Tnpo1 and Tnpo2 depletion in mouse embryonic fibroblast Wnt reporter cell lines.

**Figure 6-figure supplement 1.**
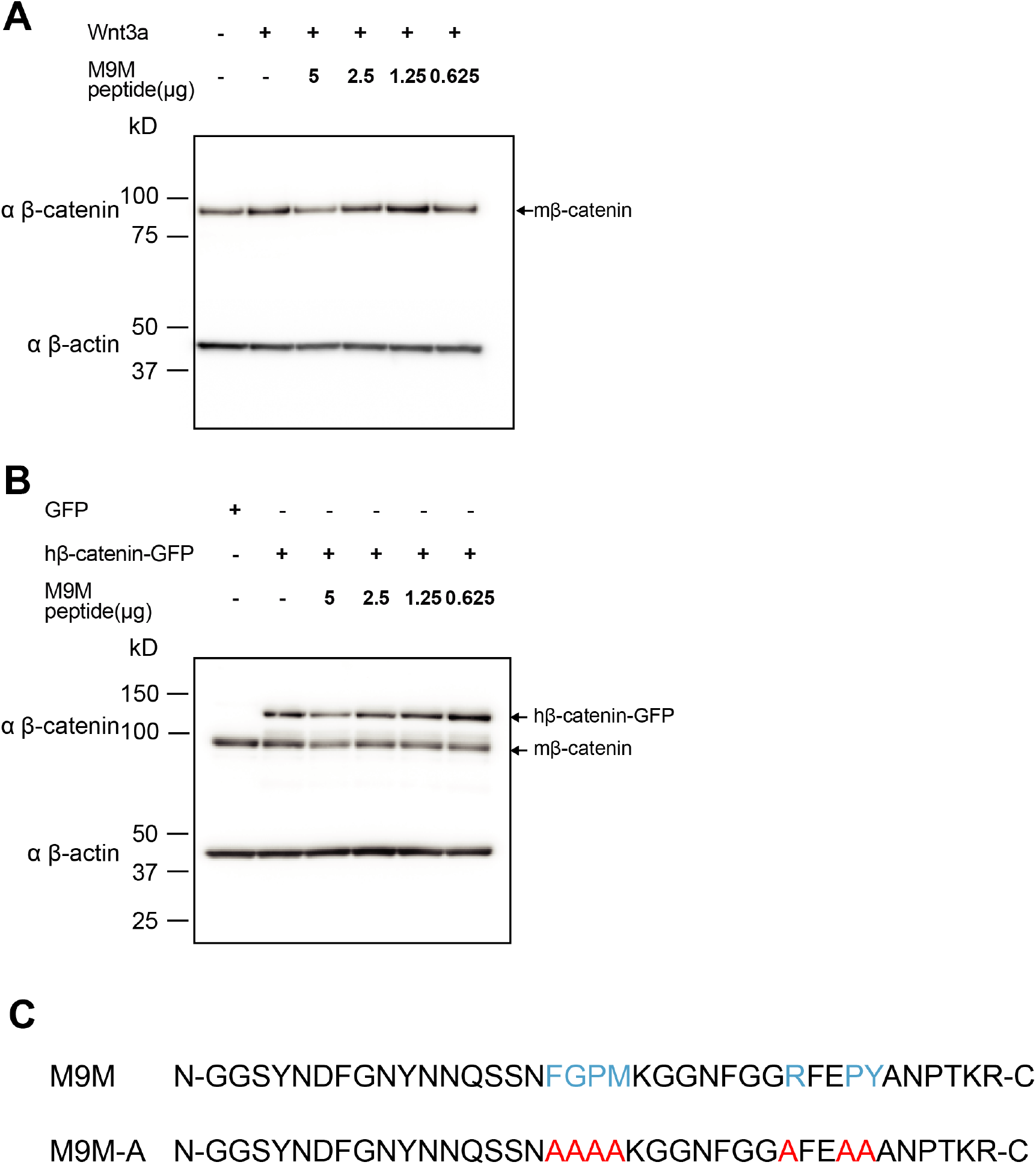
Western blots of β-catenin from 3T3 TCF/LEF luciferase assays with M9M peptide treatment. Western blot data for M9M peptide treatment in mouse embryonic fibroblast Wnt reporter cell lines. Wnt signaling is activated by **(A)** Wnt3a or **(B)** human β-catenin. **(C)** PY-NLS residues (blue) in M9M peptide were mutated to alanine (red) to create M9M-A peptide.

**Figure S8.**
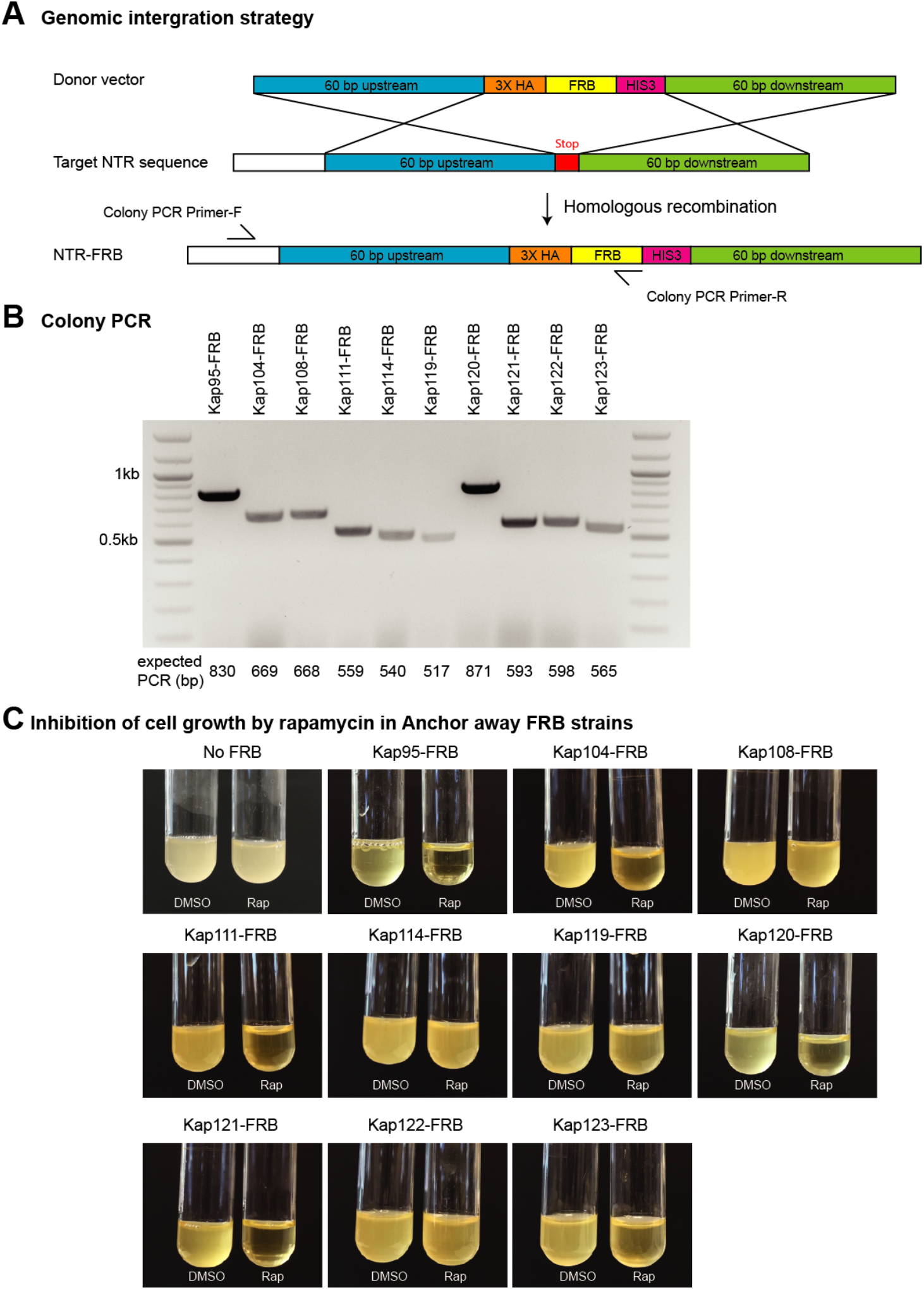
Anchor Away cloning strategy in *S. cerevisiae*. **(A)** Schematic diagram of FRB tagging to individual endogenous NTRs by homologous recombination **(B)** Screening of NTR-FRB strains by colony PCR. Primers are listed in Table S3. **(C)** Positive NTR-FRB strains from the colony PCR in **(B)** were further tested for cell growth as some NTRs are essential for survival. No FRB and DMSO were used a negative controls.

## REFERENCES

Amberg, D. C. B. D. J.; Strathern, J. N. (2005). Methods in Yeast Genetics: A Cold Spring Harbor Laboratory Course Manual. CSHL Press.

Andrade, M. A., Petosa, C., O’Donoghue, S. I., Muller, C. W., & Bork, P. (2001). Comparison of ARM and HEAT protein repeats. J Mol Biol, 309 (1), 1–18. doi:10.1006/jmbi.2001.4624

Behrens, J., von Kries, J. P., Kuhl, M., Bruhn, L., Wedlich, D., Grosschedl, R., & Birchmeier, W. (1996). Functional interaction of beta-catenin with the transcription factor LEF-1. Nature, 382 (6592), 638–642. doi:10.1038/382638a0

Bhattacharya, D., Marfo, C. A., Li, D., Lane, M., & Khokha, M. K. (2015). CRISPR/Cas9: An inexpensive, efficient loss of function tool to screen human disease genes in Xenopus. Dev Biol, 408 (2), 196–204. doi:10.1016/j.ydbio.2015.11.003

Borday, C., Parain, K., Thi Tran, H., Vleminckx, K., Perron, M., & Monsoro-Burq, A. H. (2018). An atlas of Wnt activity during embryogenesis in Xenopus tropicalis. PLoS One, 13 (4), e0193606. doi:10.1371/journal.pone.0193606

Cancer Genome Atlas, N. (2012). Comprehensive molecular characterization of human colon and rectal cancer. Nature, 487 (7407), 330–337. doi:10.1038/nature11252

Cansizoglu, A. E., Lee, B. J., Zhang, Z. C., Fontoura, B. M., & Chook, Y. M. (2007). Structure-based design of a pathway-specific nuclear import inhibitor. Nat Struct Mol Biol, 14 (5), 452–454. doi:10.1038/nsmb1229

Cavazza, T., & Vernos, I. (2015). The RanGTP Pathway: From Nucleo-Cytoplasmic Transport to Spindle Assembly and Beyond. Front Cell Dev Biol, 3, 82. doi:10.3389/fcell.2015.00082

Clevers, H., & Nusse, R. (2012). Wnt/beta-catenin signaling and disease. Cell, 149 (6), 1192– 1205. doi:10.1016/j.cell.2012.05.012

Conti, E., & Kuriyan, J. (2000). Crystallographic analysis of the specific yet versatile recognition of distinct nuclear localization signals by karyopherin alpha. Structure, 8 (3), 329–338. doi:10.1016/s0969-2126(00)00107-6

del Viso, F., & Khokha, M. (2012). Generating diploid embryos from Xenopus tropicalis. Methods Mol Biol, 917, 33–41. doi:10.1007/978-1-61779-992-1_3

Denayer, T., Tran, H. T., & Vleminckx, K. (2008). Transgenic reporter tools tracing endogenous canonical Wnt signaling in Xenopus. Methods Mol Biol, 469, 381–400. doi:10.1007/978-1-60327-469-2_24

Dormann, D., Rodde, R., Edbauer, D., Bentmann, E., Fischer, I., Hruscha, A., … Haass, C. (2010). ALS-associated fused in sarcoma (FUS) mutations disrupt Transportin-mediated nuclear import. EMBO J, 29 (16), 2841–2857. doi:10.1038/emboj.2010.143

Eleftheriou, A., Yoshida, M., & Henderson, B. R. (2001). Nuclear export of human beta-catenin can occur independent of CRM1 and the adenomatous polyposis coli tumor suppressor. J Biol Chem, 276 (28), 25883–25888. doi:10.1074/jbc.M102656200

Fagotto, F. (2013). Looking beyond the Wnt pathway for the deep nature of beta-catenin. EMBO Rep, 14 (5), 422–433. doi:10.1038/embor.2013.45

Fagotto, F., Gluck, U., & Gumbiner, B. M. (1998). Nuclear localization signal-independent and importin/karyopherin-independent nuclear import of beta-catenin. Curr Biol, 8 (4), 181– 190. doi:10.1016/s0960-9822(98)70082-x

Fontes, M. R., Teh, T., & Kobe, B. (2000). Structural basis of recognition of monopartite and bipartite nuclear localization sequences by mammalian importin-alpha. J Mol Biol, 297 (5), 1183–1194. doi:10.1006/jmbi.2000.3642

Goto, T., Sato, A., Adachi, S., Iemura, S., Natsume, T., & Shibuya, H. (2013). IQGAP1 protein regulates nuclear localization of beta-catenin via importin-beta5 protein in Wnt signaling. J Biol Chem, 288 (51), 36351–36360. doi:10.1074/jbc.M113.520528

Griffin, J. N., Del Viso, F., Duncan, A. R., Robson, A., Hwang, W., Kulkarni, S., … Khokha, M. K. (2018). RAPGEF5 Regulates Nuclear Translocation of beta-Catenin. Dev Cell, 44 (2), 248–260 e244. doi:10.1016/j.devcel.2017.12.001

Haruki, H., Nishikawa, J., & Laemmli, U. K. (2008). The anchor-away technique: rapid, conditional establishment of yeast mutant phenotypes. Mol Cell, 31 (6), 925–932. doi:10.1016/j.molcel.2008.07.020

Heasman, J., Crawford, A., Goldstone, K., Garner-Hamrick, P., Gumbiner, B., McCrea, P., … Wylie, C. (1994). Overexpression of cadherins and underexpression of beta-catenin inhibit dorsal mesoderm induction in early Xenopus embryos. Cell, 79 (5), 791–803. doi:10.1016/0092-8674(94)90069-8

Heasman, J., Kofron, M., & Wylie, C. (2000). Beta-catenin signaling activity dissected in the early Xenopus embryo: a novel antisense approach. Dev Biol, 222 (1), 124–134. doi:10.1006/dbio.2000.9720

Henderson, B. R. (2000). Nuclear-cytoplasmic shuttling of APC regulates beta-catenin subcellular localization and turnover. Nat Cell Biol, 2 (9), 653–660. doi:10.1038/35023605

Holstein, T. W. (2012). The evolution of the Wnt pathway. Cold Spring Harb Perspect Biol, 4 (7), a007922. doi:10.1101/cshperspect.a007922

Huber, A. H., Nelson, W. J., & Weis, W. I. (1997). Three-dimensional structure of the armadillo repeat region of beta-catenin. Cell, 90 (5), 871–882. doi:10.1016/s0092-8674(00)80352-9

Huber, O., Korn, R., McLaughlin, J., Ohsugi, M., Herrmann, B. G., & Kemler, R. (1996). Nuclear localization of beta-catenin by interaction with transcription factor LEF-1. Mech Dev, 59 (1), 3–10. doi:10.1016/0925-4773(96)00597-7

Kadowaki, T., Zhao, Y., & Tartakoff, A. M. (1992). A conditional yeast mutant deficient in mRNA transport from nucleus to cytoplasm. Proc Natl Acad Sci U S A, 89 (6), 2312– 2316. doi:10.1073/pnas.89.6.2312

Khokha, M. K., Chung, C., Bustamante, E. L., Gaw, L. W., Trott, K. A., Yeh, J., … Grammer, T. C. (2002). Techniques and probes for the study of Xenopus tropicalis development. Dev Dyn, 225 (4), 499–510. doi:10.1002/dvdy.10184

Khokha, M. K., Yeh, J., Grammer, T. C., & Harland, R. M. (2005). Depletion of three BMP antagonists from Spemann’s organizer leads to a catastrophic loss of dorsal structures. Dev Cell, 8 (3), 401–411. doi:10.1016/j.devcel.2005.01.013

Koike, M., Kose, S., Furuta, M., Taniguchi, N., Yokoya, F., Yoneda, Y., & Imamoto, N. (2004). beta-Catenin shows an overlapping sequence requirement but distinct molecular interactions for its bidirectional passage through nuclear pores. J Biol Chem, 279 (32), 34038–34047. doi:10.1074/jbc.M405821200

Komiya, Y., Mandrekar, N., Sato, A., Dawid, I. B., & Habas, R. (2014). Custos controls beta-catenin to regulate head development during vertebrate embryogenesis. Proc Natl Acad Sci U S A, 111 (36), 13099–13104. doi:10.1073/pnas.1414437111

Kosugi, S., Hasebe, M., Tomita, M., & Yanagawa, H. (2008). Nuclear export signal consensus sequences defined using a localization-based yeast selection system. Traffic, 9 (12), 2053– 2062. doi:10.1111/j.1600-0854.2008.00825.x

Lange, A., Mills, R. E., Devine, S. E., & Corbett, A. H. (2008). A PY-NLS nuclear targeting signal is required for nuclear localization and function of the Saccharomyces cerevisiae mRNA-binding protein Hrp1. J Biol Chem, 283 (19), 12926–12934. doi:10.1074/jbc.M800898200

Lee, B. J., Cansizoglu, A. E., Suel, K. E., Louis, T. H., Zhang, Z., & Chook, Y. M. (2006). Rules for nuclear localization sequence recognition by karyopherin beta 2. Cell, 126 (3), 543–558. doi:10.1016/j.cell.2006.05.049

Longtine, M. S., McKenzie, A., 3rd, Demarini, D. J., Shah, N. G., Wach, A., Brachat, A., … Pringle, J. R. (1998). Additional modules for versatile and economical PCR-based gene deletion and modification in Saccharomyces cerevisiae. Yeast, 14 (10), 953–961. doi:10.1002/(SICI)1097-0061(199807)14:10<953::AID-YEA293>3.0.CO;2-U

MacDonald, B. T., Tamai, K., & He, X. (2009). Wnt/beta-catenin signaling: components, mechanisms, and diseases. Dev Cell, 17 (1), 9–26. doi:10.1016/j.devcel.2009.06.016

Malik, H. S., Eickbush, T. H., & Goldfarb, D. S. (1997). Evolutionary specialization of the nuclear targeting apparatus. Proc Natl Acad Sci U S A, 94 (25), 13738–13742. doi:10.1073/pnas.94.25.13738

McMahon, A. P., & Moon, R. T. (1989). Ectopic expression of the proto-oncogene int-1 in Xenopus embryos leads to duplication of the embryonic axis. Cell, 58 (6), 1075–1084. doi:10.1016/0092-8674(89)90506-0

Mis, M., O’Brien, S., Steinhart, Z., Lin, S., Hart, T., Moffat, J., & Angers, S. (2020). IPO11 mediates betacatenin nuclear import in a subset of colorectal cancers. J Cell Biol, 219(2). doi:10.1083/jcb.201903017

Molenaar, M., van de Wetering, M., Oosterwegel, M., Peterson-Maduro, J., Godsave, S., Korinek, V., … Clevers, H. (1996). XTcf-3 transcription factor mediates beta-catenin-induced axis formation in Xenopus embryos. Cell, 86 (3), 391–399. doi:10.1016/s0092-8674(00)80112-9

Moon, R. T., Brown, J. D., & Torres, M. (1997). WNTs modulate cell fate and behavior during vertebrate development. Trends Genet, 13 (4), 157–162. doi:10.1016/s0168-9525(97)01093-7

Moon, R. T., Kohn, A. D., De Ferrari, G. V., & Kaykas, A. (2004). WNT and beta-catenin signalling: diseases and therapies. Nat Rev Genet, 5 (9), 691–701. doi:10.1038/nrg1427

Moreno-Mateos, M. A., Vejnar, C. E., Beaudoin, J. D., Fernandez, J. P., Mis, E. K., Khokha, M. K., & Giraldez, A. J. (2015). CRISPRscan: designing highly efficient sgRNAs for CRISPR-Cas9 targeting in vivo. Nat Methods, 12 (10), 982–988. doi:10.1038/nmeth.3543

Morin, P. J., Sparks, A. B., Korinek, V., Barker, N., Clevers, H., Vogelstein, B., & Kinzler, K. W. (1997). Activation of beta-catenin-Tcf signaling in colon cancer by mutations in betacatenin or APC. Science, 275 (5307), 1787–1790. doi:10.1126/science.275.5307.1787

Neufeld, K. L., Zhang, F., Cullen, B. R., & White, R. L. (2000). APC-mediated downregulation of beta-catenin activity involves nuclear sequestration and nuclear export. EMBO Rep, 1 (6), 519–523. doi:10.1093/embo-reports/kvd117

Niehrs, C. (2012). The complex world of WNT receptor signalling. Nat Rev Mol Cell Biol, 13 (12), 767–779. doi:10.1038/nrm3470

Nishisho, I., Nakamura, Y., Miyoshi, Y., Miki, Y., Ando, H., Horii, A., … Hedge, P. (1991). Mutations of chromosome 5q21 genes in FAP and colorectal cancer patients. Science, 253 (5020), 665–669. doi:10.1126/science.1651563

Nusse, R., & Varmus, H. E. (1982). Many tumors induced by the mouse mammary tumor virus contain a provirus integrated in the same region of the host genome. Cell, 31 (1), 99–109. doi:10.1016/0092-8674(82)90409-3

Orsulic, S., & Peifer, M. (1996). An in vivo structure-function study of armadillo, the beta-catenin homologue, reveals both separate and overlapping regions of the protein required for cell adhesion and for wingless signaling. J Cell Biol, 134 (5), 1283–1300. doi:10.1083/jcb.134.5.1283

Polakis, P. (2012). Wnt signaling in cancer. Cold Spring Harb Perspect Biol, 4(5). doi:10.1101/cshperspect.a008052

Popken, P., Ghavami, A., Onck, P. R., Poolman, B., & Veenhoff, L. M. (2015). Size-dependent leak of soluble and membrane proteins through the yeast nuclear pore complex. Mol Biol Cell, 26 (7), 1386–1394. doi:10.1091/mbc.E14-07-1175

Prieve, M. G., & Waterman, M. L. (1999). Nuclear localization and formation of beta-catenin- lymphoid enhancer factor 1 complexes are not sufficient for activation of gene expression. Mol Cell Biol, 19 (6), 4503–4515. doi:10.1128/mcb.19.6.4503

Rebane, A., Aab, A., & Steitz, J. A. (2004). Transportins 1 and 2 are redundant nuclear import factors for hnRNP A1 and HuR. RNA, 10 (4), 590–599. doi:10.1261/rna.5224304

Rosin-Arbesfeld, R., Townsley, F., & Bienz, M. (2000). The APC tumour suppressor has a nuclear export function. Nature, 406 (6799), 1009–1012. doi:10.1038/35023016

Schmidt, H. B., & Gorlich, D. (2016). Transport Selectivity of Nuclear Pores, Phase Separation, and Membraneless Organelles. Trends Biochem Sci, 41 (1), 46–61. doi:10.1016/j.tibs.2015.11.001

Sempou, E., Lakhani, O. A., Amalraj, S., & Khokha, M. K. (2018). Candidate Heterotaxy Gene FGFR4 Is Essential for Patterning of the Left-Right Organizer in Xenopus. Front Physiol, 9, 1705. doi:10.3389/fphys.2018.01705

Sharma, M., Jamieson, C., Lui, C., & Henderson, B. R. (2014). WITHDRAWN: The hydrophobic rich N- and C-terminal tails of beta-catenin facilitate nuclear import of beta-catenin. J Biol Chem. doi:10.1074/jbc.M114.603209

Sharma, M., Johnson, M., Brocardo, M., Jamieson, C., & Henderson, B. R. (2014). Wnt signaling proteins associate with the nuclear pore complex: implications for cancer. Adv Exp Med Biol, 773, 353–372. doi:10.1007/978-1-4899-8032-8_16

Sive, H. L., Grainger, R. M., & Harland, R. M. (2007). Xenopus laevis In Vitro Fertilization and Natural Mating Methods. CSH Protoc, 2007, pdb prot4737. doi:10.1101/pdb.prot4737

Smith, W. C., & Harland, R. M. (1991). Injected Xwnt-8 RNA acts early in Xenopus embryos to promote formation of a vegetal dorsalizing center. Cell, 67 (4), 753–765. doi:10.1016/0092-8674(91)90070-f

Sokol, S., Christian, J. L., Moon, R. T., & Melton, D. A. (1991). Injected Wnt RNA induces a complete body axis in Xenopus embryos. Cell, 67 (4), 741–752. doi:10.1016/0092-8674(91)90069-b

Soniat, M., & Chook, Y. M. (2015). Nuclear localization signals for four distinct karyopherin-beta nuclear import systems. Biochem J, 468 (3), 353–362. doi:10.1042/BJ20150368

Soniat, M., Sampathkumar, P., Collett, G., Gizzi, A. S., Banu, R. N., Bhosle, R. C., … Almo, S. C. (2013). Crystal structure of human Karyopherin beta2 bound to the PY-NLS of Saccharomyces cerevisiae Nab2. J Struct Funct Genomics, 14 (2), 31–35. doi:10.1007/s10969-013-9150-1

Suh, E. K., & Gumbiner, B. M. (2003). Translocation of beta-catenin into the nucleus independent of interactions with FG-rich nucleoporins. Exp Cell Res, 290 (2), 447–456. doi:10.1016/s0014-4827(03)00370-7

Timney, B. L., Raveh, B., Mironska, R., Trivedi, J. M., Kim, S. J., Russel, D., … Rout, M. P. (2016). Simple rules for passive diffusion through the nuclear pore complex. J Cell Biol, 215 (1), 57–76. doi:10.1083/jcb.201601004

Twyffels, L., Gueydan, C., & Kruys, V. (2014). Transportin-1 and Transportin-2: protein nuclear import and beyond. FEBS Lett, 588 (10), 1857–1868. doi:10.1016/j.febslet.2014.04.023

van de Wetering, M., Cavallo, R., Dooijes, D., van Beest, M., van Es, J., Loureiro, J., … Clevers, H. (1997). Armadillo coactivates transcription driven by the product of the Drosophila segment polarity gene dTCF. Cell, 88 (6), 789–799. doi:10.1016/s0092-8674(00)81925-x

van de Wetering, M., Oosterwegel, M., Dooijes, D., & Clevers, H. (1991). Identification and cloning of TCF-1, a T lymphocyte-specific transcription factor containing a sequence-specific HMG box. EMBO J, 10 (1), 123-132. Retrieved from https://www.ncbi.nlm.nih.gov/pubmed/1989880

Waterhouse, A. M., Procter, J. B., Martin, D. M., Clamp, M., & Barton, G. J. (2009). Jalview Version 2--a multiple sequence alignment editor and analysis workbench. Bioinformatics, 25 (9), 1189–1191. doi:10.1093/bioinformatics/btp033

Weis, K. (2003). Regulating access to the genome: nucleocytoplasmic transport throughout the cell cycle. Cell, 112 (4), 441–451. doi:10.1016/s0092-8674(03)00082-5

Wente, S. R., & Rout, M. P. (2010). The nuclear pore complex and nuclear transport. Cold Spring Harb Perspect Biol, 2 (10), a000562. doi:10.1101/cshperspect.a000562

Wood, L. D., Parsons, D. W., Jones, S., Lin, J., Sjoblom, T., Leary, R. J., … Vogelstein, B. (2007). The genomic landscapes of human breast and colorectal cancers. Science, 318 (5853), 1108–1113. doi:10.1126/science.1145720

Wozniak, R. W., Rout, M. P., & Aitchison, J. D. (1998). Karyopherins and kissing cousins. Trends Cell Biol, 8 (5), 184–188. doi:10.1016/s0962-8924(98)01248-3

Xu, L., & Massague, J. (2004). Nucleocytoplasmic shuttling of signal transducers. Nat Rev Mol Cell Biol, 5 (3), 209–219. doi:10.1038/nrm1331

Yano, R., Oakes, M. L., Tabb, M. M., & Nomura, M. (1994). Yeast Srp1p has homology to armadillo/plakoglobin/beta-catenin and participates in apparently multiple nuclear functions including the maintenance of the nucleolar structure. Proc Natl Acad Sci U S A, 91 (15), 6880–6884. doi:10.1073/pnas.91.15.6880

Yokoya, F., Imamoto, N., Tachibana, T., & Yoneda, Y. (1999). beta-catenin can be transported into the nucleus in a Ran-unassisted manner. Mol Biol Cell, 10 (4), 1119–1131. doi:10.1091/mbc.10.4.1119

